# Identification of immunogenic and cross-reactive chikungunya virus-specific CD4^+^ T cell epitopes in chronic chikungunya viral arthritic disease in humans

**DOI:** 10.1101/2024.10.22.619717

**Authors:** Rimjhim Agarwal, Calvin Ha, Fernanda H. Cortes, Yeji Lee, Amparo Martínez-Pérez, Rosa Isela Gálvez, Izabella N. Castillo, Angel Balmaseda, Eva Harris, Claudia M Romero-Vivas, Lakshmanane Premkumar, Andrew K Falconar, Alba Grifoni, Alessandro Sette, Daniela Weiskopf

**Affiliations:** Center for Vaccine Innovation, La Jolla Institute for Immunology (LJI), La Jolla, CA 92037, USA; Biomedical Sciences Graduate Program, School of Medicine, University of California San Diego (UCSD), La Jolla, CA 92037, USA; Laboratory of AIDS and Molecular Immunology, Institute Oswaldo Cruz, Fiocruz, Rio de Janeiro, RJ 21040-360, Brazil; Department of Microbiology and Immunology, University of North Carolina School of Medicine, Chapel Hill, NC 27599, USA; Sustainable Sciences Institute, Managua 14006, Nicaragua; Division of Infectious Diseases and Vaccinology, School of Public Health, University of California, Berkeley, Berkeley, CA 94720-3370, USA; Laboratorio de Enfermedades Tropicales, Departamento de Medicina, Fundación Universidad del Norte, Barranquilla, Colombia; Department of Medicine, Division of Infectious Diseases and Global Public Health, University of California San Diego (UCSD), La Jolla, CA 92037, USA

**Keywords:** Chikungunya virus, chronic viral disease, CD4^+^ T cells, T cell epitopes, alphaviruses, conservation

## Abstract

Chikungunya virus (CHIKV), a mosquito-borne alphavirus, causes acute febrile illness that can progress into chronic chikungunya virus disease (CHIKVD) marked by persistent debilitating arthralgia. At present, the exact cause of chronic CHIKVD is not understood, and in humans, the targets of CD4^+^ T cells in CHIKV are currently unknown. Here, by stimulating peripheral blood mononuclear cells (PBMCs) collected from patients suffering from chronic CHIKVD with peptides spanning the entire CHIKV genome, we provide a comprehensive landscape of CHIKV CD4^+^ T cell epitopes. We identified 123 novel CD4^+^ T cell epitopes and three immunodominant regions in E1, nsP1 and CP proteins. The immunodominance of these E1, nsP1 and CP regions was mapped to optimal epitopes, characterized by the capacity to bind to many common HLA class II allelic variants. In addition, we designed and validated a new CHIKV-specific CD4^+^ T cell epitope megapool, spanning both structural and non-structural proteins, which can be a useful tool to study CHIKV-specific T cell responses in small blood volumes, typically available in pediatric or clinical samples. Finally, by *in silico* assessment of the conservation of the CHIKV proteome in a diverse set of alphaviruses, we defined CHIKV epitopes conserved across arthritogenic and encephalitic viruses. Overall, our work is the first to identify CD4^+^ T cell targets of CHIKV in humans, expanding our capacity to study the role of T cells in CHIKV pathogenesis and mapping targets of alphaviruses for vaccine design.

## INTRODUCTION

Alphaviruses are arthropod-borne viruses from the *Togaviridae* family that cause major epidemics around the globe. The *Alphavirus* genus is categorized into two groups based on genetic relatedness and symptomatology. Arthritogenic alphaviruses (predominantly Old-World viruses), such as chikungunya virus (CHIKV), O’nyong-nyong virus (ONNV) and Ross River virus (RRV), cause persistent rheumatic symptoms while encephalitic alphaviruses (predominantly New World viruses), such as Venezuelan equine encephalitic virus (VEEV) and eastern equine encephalitic virus (EEEV), lead to incapacitating neurological manifestations^1–4^. Although CHIKV is generally considered an arthritogenic virus, a few cases of encephalitis have been reported in neonates infected with CHIKV during recent epidemics^5^.

CHIKV, a prototypic arthritogenic alphavirus, is the causative agent for chronic chikungunya virus disease (CHIKVD) which is characterized by persistent debilitating arthralgia in small joints in 30-60% of infected individuals^6–9^. The virus primarily circulates in tropical and subtropical regions, including South/Southeast Asia, Africa, and Central and South America^10^. The high morbidity rate and recurrent outbreaks due to the expanding range of *Aedes* mosquitoes pose a significant global public health threat^11,12^.

The CHIKV genome is 12kb in length and consists of four non-structural proteins (nsP1, nsP2, nsP3 and nsP4) and five structural proteins (Capsid, Envelope 1-3 [E1-3] and 6K)^13^. Non-structural proteins play a pivotal role in viral replication and structural proteins encode for proteins required for virion assembly^14^. IXCHIQ, the first FDA-approved CHIKV vaccine, consists of both non-structural and structural proteins with large deletions in the nsP3 protein^15^.

Currently, the cause of chronic arthralgia post-CHIKV infection is poorly understood; however, numerous studies have alluded to the inflammatory role of CD4^+^ T cells. Previous studies in mice have shown extensive infiltration of CD4^+^ T cells in inflamed joints of infected mice^16,17^. In addition, the adoptive transfer of CD4^+^ T cells from CHIKV-infected mice into CHIKV-infected T cell receptor knock-out (TCR^-/-^) mice led to severe joint swelling, vascular leakage and cellular infiltration^18^. Our recent unpublished work has shown a significantly higher frequency of CHIKV-specific CD4^+^ T cells and negligible CD8^+^ T cell responses in the peripheral blood of patients with chronic CHIKVD^19^, consistent with an association of CHIKV-specific CD4^+^ T cells with chronic viral arthritic disease.

However, the epitope targets of CD4^+^ T cell responses in chronic CHIKVD in humans are currently unknown, thereby hindering our further understanding of the role of CD4^+^ T cells in chronic CHIKVD and the development of effective vaccines. Few studies have experimentally described CD4^+^ T cell epitopes in mice during the acute phase of infection, where dominant epitopes were found in the nsP1, E1 and E2 proteins^18,20^. Specifically, Teo et al. discovered two epitopes in nsP1_145-162_ and E2_2800-2818_ proteins, which when transferred as epitope-specific T cell lines into TCR^-/-^ mice induced joint inflammation^18^. While several groups have developed methods to predict T cell epitopes *in silico* ^21–23^, to our knowledge, no studies have experimentally analyzed specific T cell epitope sequences in humans suffering from chronic CHIKVD or other alphavirus infections in the context of natural infection.

In this study, by *in vitro* stimulation of peripheral blood mononuclear cells (PBMCs) collected from individuals affected by chronic CHIKVD with peptides spanning the entire CHIKV proteome, we map the landscape of human CD4^+^ T cell epitopes identified during the chronic phase of CHIKVD. We describe123 novel CD4^+^ T cell epitopes of CHIKV and identify immunodominant regions and epitopes in the E1, nsP1 and CP proteins. Additionally, we find a high degree of conservation of the identified CHIKV epitopes in other arthritogenic alphaviruses and to a lower degree in encephalitic alphaviruses. Overall, our work comprehensively characterizes the immunogenic targets of circulating CHIKV-specific CD4^+^ T cells in the periphery in humans, expanding our understanding of the pathogenesis of alphaviruses, which is essential to develop and characterize effective vaccines.

## RESULTS

### Experimental design for proteome-wide screen of CHIKV human T cell epitopes

We designed a comprehensive peptide library to map the T cell epitopes associated with CHIKV infection in humans. Following the previously described megapool approach (MP)^19,24^, we synthesized 15-mer overlapping peptides for four non-structural CHIKV proteins (nsP1, nsP2, nsP3, nsP4) and five structural CHIKV proteins (Capsid or CP, E3, E2, 6K, E1) and additional frequent variant peptides as described in the Methods section (Figure 1A). Overall, we synthesized a total of 992 peptides, which comprised 741 consensus peptides and 251 variant peptides. These peptides were organized in ten MPs, which were also further divided into smaller Mesopools (MS) of 9-10 individual peptides.

**Figure 1:**
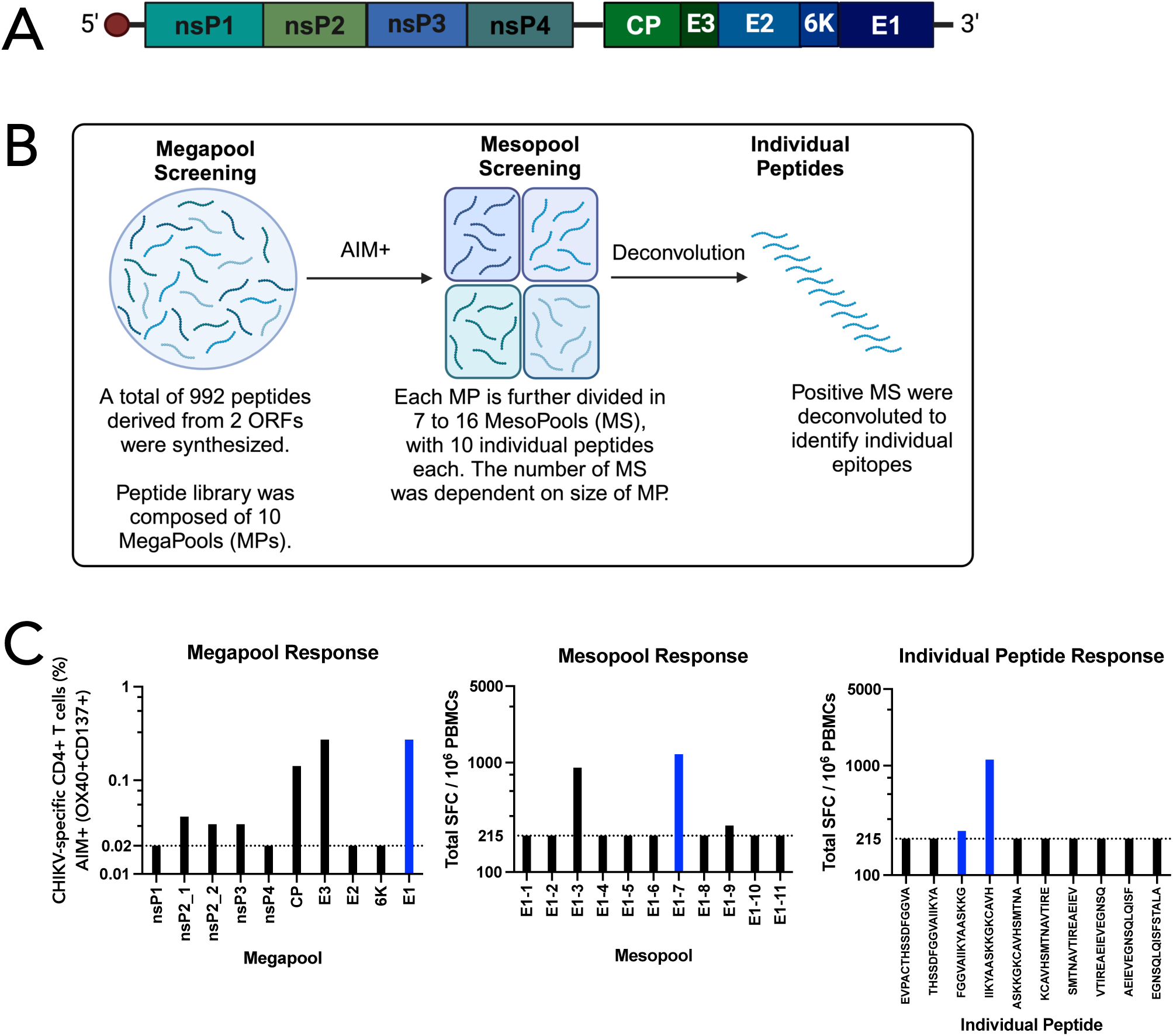
Experimental workflow for screening CD4^+^ T cell epitopes in CHIKV. **(A)** Schematic representation of CHIKV proteome comprising four non-structural (nsP1, nsP2, nsP3 and nsP4) and five structural proteins (Capsid or CP, E3, E2 and E1). **(B)** Workflow of epitope screening. All donors were tested in the AIM assay by stimulation with 10 megapools (MP) corresponding to each CHIKV protein (nsP1, nsP2, nsP3, nsP4, CP, E3, E2, 6K, E1). Positive donors in the AIM assay (OX40+CD137+ or OX40+CD40L+) were tested in the FluoroSpot assay by stimulating with smaller pools of MP, called mesopools (MS), each of which contained 9-10 individual peptides. Each MS was deconvoluted to determine individual epitopes. **(C)** An example of the experimental workflow for the responses of one donor to the E1 protein. The first panel shows responses in the AIM assay to all CHIKV MPs. The middle panel shows SFCs per million PBMCs to each MS of the E1 protein. The third panel depicts responses to individual peptides in E1-7 MS. The dotted line shows the threshold of positivity. The blue highlighted bars depict an example of a positive response from one donor.

The library was used to map T cell epitopes recognized in 17 individuals from Colombia who displayed chronic arthritis-like symptoms on average 6.3 years post-CHIKV infection (Table 1). Infections were confirmed by measuring CHIKV-specific IgG titers (Figure S1A). To identify the global pattern of recognition, PBMCs from each donor were assayed *ex vivo* with each CHIKV protein MP (nsP1, nsP2_1, nsP2_2, nsP3, nsP4, CP, E3, E2, 6K, E1) (Figure S1C). As described before^19^, negligible CD8^+^ T cell responses were detected; hence, we focused our epitope identification efforts on CD4^+^ T cells. CHIKV-specific CD4^+^ T cells were identified in an activation-induced marker assay (AIM assay), using the upregulation of either OX40+CD137+ or OX40+CD40L+ markers, both of which are commonly used to measure antigen-specific CD4^+^ T cell responses^25–27^. Using two different AIM markers allowed us to perform a more comprehensive screening to identify more positive responses. As previously described, responses greater than the limit of sensitivity (LOS) and stimulation index (S.I.) greater than two were considered positive. CHIKV-specific CD4^+^ T cell responses were detected in 88% (15/17) of donors. The highest frequency of AIM+ OX40+CD137+ responses were detected against E1 (65%), E2 (53%), CP (29%) and nsP3 (29%) proteins, with a similar trend for AIM+ OX40+CD40L+ responses, where E1 (59%), E2 (53%), CP (41%) and nsP1 (41%) proteins were most frequently recognized (Figure S1C). Of 170 unique donor-protein combinations tested, a total of 47 positive responses were detected marked by upregulation of OX40+CD137+ and 57 positive responses were detected marked by upregulation of OX40+CD40L+ (Figure S1C).

**Table 1:** Characteristics of the donor cohort.

| Characteristic | CHIKV seronegative cohort | Epitope Screening cohort | Fine epitope mapping cohort |  | Epitope MP Validation cohort |  |
| --- | --- | --- | --- | --- | --- | --- |
|  |  |  | Chronic | Recovered | Chronic | Uninfected |
| Donors, n | 6 | 17 | 10 | 4 | 19 | 7 |
| Gender, n (%) |  |  |  |  |  |  |
| Male | NA | 6 (35%) | 2 (20%) | 0 (0%) | 6 (32%) | 2 (29%) |
| Female | NA | 11 (65%) | 8 (80%) | 4 (100%) | 13 (68%) | 5 (71%) |
| Age *, years (mean ± SD) | NA | 46.8 ± 14 | 44.1 ± 13 | 34.5 ± 2.6 | 43.9 ± 16 | 43.9 ± 12 |
| Years post-infection (mean ± SD) |  | 6.3 ± 0.5 | 6.2 ± 0.6 | 6.4 ± 0.3 | 6.3 ± 0.4 |  |
| Current symptoms |  |  |  |  |  |  |
| Arthralgia** (%) | 0 | 100 | 100 | 0 | 100 | 0 |
| CHIKV serostatus | Negative | Positive | Positive | Positive | Positive | Negative |
| Site | Nicaragua | Colombia | Colombia |  | Colombia |  |

\* At the time of sample collection
\*\* Includes pain in joints and upper and lower limbs, swelling in limbs, and muscle pain
NA - Information not available

### Identification of CHIKV human CD4^+^ T cell epitopes

Donor/MP combinations that were positive in the AIM assay were restimulated *in vitro* to allow for epitope identification. In case of limiting cell availability, the combinations with the strongest AIM+ responses were selected for re-stimulation. After 14 days, each T cell culture was tested against the MS corresponding to the stimulating MP in a FluoroSpot assay. The threshold of positivity (215 spot-forming cells per million PBMCs or SFC/10^6^ PBMCs) for the FluoroSpot assay was determined as the value higher than 99% of all the measured responses following the same *in vitro* stimulation protocol of PBMCs from six CHIKV-seronegative donors from Nicaragua with all CHIKV mesopools (Table 1 and Figure S1D).

Overall, of a total of 795 MS tested, 293 (37%) MS were associated with positive responses. Cultures from these positive MS responses were further tested with individual peptides to identify specific epitopes. As a result, 123 consensus epitopes and 24 variant epitopes were identified (Table S1). A summary of the screening strategy is shown in Figure 1B and representative data from one donor is shown in Figure 1C. The representative donor recognized six MPs (nsP2_1, nsP2_2, nsP3, CP, E3 and E1) in the AIM assay, which were then screened with MS. From the E1 MS, the donor recognized three MS (E1-3, E1-7 and E1-9). The E1-7 MS, among other pools, was then deconvoluted where two epitopes from the pool were recognized by the donor. As the variant peptides overlap with the consensus peptides, we chose to focus on consensus epitopes for further analysis.

### Immunodominant proteins in CHIKV proteome recognized by CD4^+^ T cells

Post-deconvolution, we identified a total of 123 epitopes from the consensus genome (Figure 2A). These sequences have been submitted to IEDB (submission IDs 1000910, 1000911 and 1000913). The majority of these epitopes were from E1 (24%), nsP1 (21%) and CP (16%) proteins with few epitopes located in nsP4 (2%) and 6K (2%), while none were identified in the E3 protein. Overall, non-structural and structural proteins accounted for 34% and 66% of total magnitude of the response, respectively, with the highest total magnitude of responses against epitopes in E1 (35%), nsP1 (21%) and CP (18%) (Figure 2B). Notably, in addition to having the highest number of epitopes and the highest frequency of response, the E1 protein elicited over one-third of the overall magnitude of the response.

**Figure 2:**
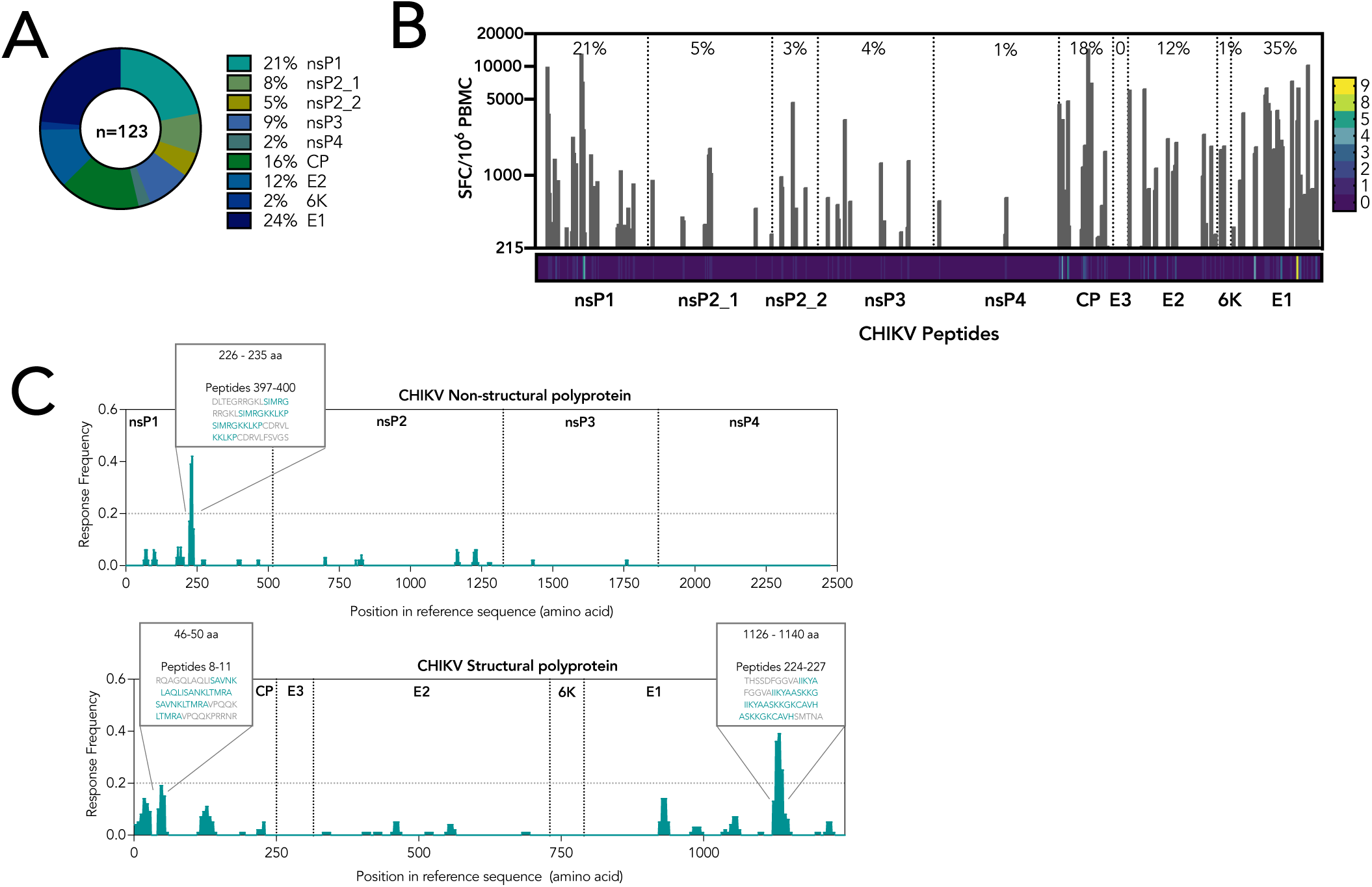
Immunodominant proteins in CHIKV proteome recognized by CD4^+^ T cells. **(A)** Frequency (%) of epitopes detected per CHIKV protein. n refers to the number of total epitopes identified. **(B)** The magnitude (SFC/10^6^ PBMCs) of positive response to each epitope in the entire CHIKV proteome. The percentage shows the percent of overall magnitude of responses elicited by each protein. The heatmap under the graph indicate the number of donors who recognize each epitope, ranging from 0-9 donors. **(C)** The lower 95% confidence interval of response frequency of each epitope plotted for the non-structural (top) and structural polyprotein (bottom). The dotted line indicates the threshold of positivity (0.2). The highlighted regions depict the residue number that reach the threshold of positivity and the peptide sequences.

To identify the immunodominant regions in CHIKV proteome, we visualized the identified epitopes to the CHIKV proteome using the standalone ImmunomeBrowser tool from the Immune Epitope Database Analysis Resource (IEDB-AR)^28^ and calculated the lower 95% confidence interval of the response frequency of each peptide in structural and non-structural polyproteins separately. Two regions - nsP1_226-235_ and E1_1126-1140_ – exceeded a threshold of positivity of 20%, while CP_46-50_ reached the threshold (Figure 2C).

### Immunodominant epitopes in CHIKV proteome recognized by CD4+ T cells

Next, we focused our analysis on responses to individual epitopes. The magnitude of response from epitopes varied widely, ranging from 200 to 21,200 SFCs/10^6^ PBMCs (average response was 1128 SFCs/10^6^ PBMCs; Figure 3A). Interestingly, 38 epitopes accounted for 75% of the total magnitude of interferon gamma (IFN*γ*) response from all donors tested (Figure 3B and Table 2).

**Figure 3:**
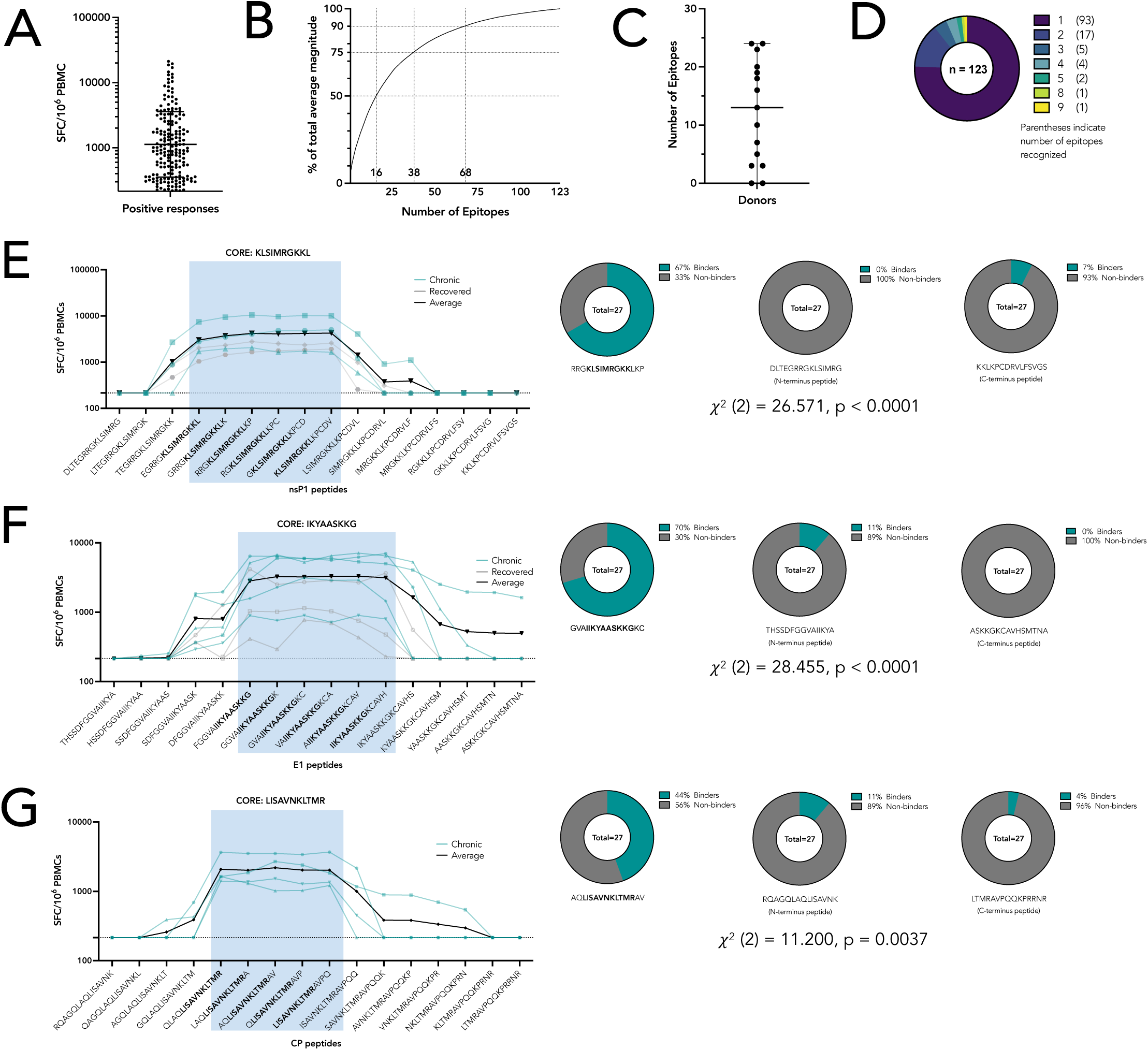
Immunodominant epitopes in CHIKV recognized by CD4^+^ T cells. **(A)** The magnitude of all positive responses, shown as the total number of IFN*γ* spot-forming cells per million PBMCs (SFC/10^6^ PBMCs). Average response is at 1128 SFC/10^6^ PBMCs with responses varying from 220-21200 SFC/10^6^ PBMCs. Each dot refers to a CHIKV epitope**. (B)** Percentage of total of average response for all donors plotted as the function of the total number of epitopes. Epitopes were ranked in descending order of average magnitude and percentage of total magnitude was calculated based on cumulative sum of response. Dotted lines represent the number of epitopes that account for 50%, 75% and 90% of total responses. **(C)** Number of epitopes recognized by each donor. An average of 12 epitopes were identified. **(D)** Pie chart indicates the frequency of response for each epitope. Parenthesis indicates the number of epitopes recognized by the specified number of donors. **(E), (F) and (G)** SFC/10^6^ PBMCs shown for individual chronic (blue line) and recovered (gray line) donors against 15mer peptides overlapping by 14 residues sequentially spanning the immunodominant region of **(E)** the nsP1 (chronic: n = 3, recovered: n = 2), **(F)** E1 (chronic: n = 5, recovered: n = 3) and **(G)** CP (chronic: n = 4) regions. The black line shows the average SFC/10^6^ PBMCs for all donors shown. Blue highlighted regions and bolded peptide sequences indicate region with the highest response and the common amino acid sequence, respectively. The pie charts on the right depict the percentage of HLA alleles predicted to bind to the peptide sequences (i.e. the mapped epitope and negative control peptides on the N- and C-termini) shown under the pie chart. Chi-square value with degree of freedom and p-value is reported for all peptide sequences. Data are represented as mean ± SD or geomean ± geometric SD.

**Table 2:**
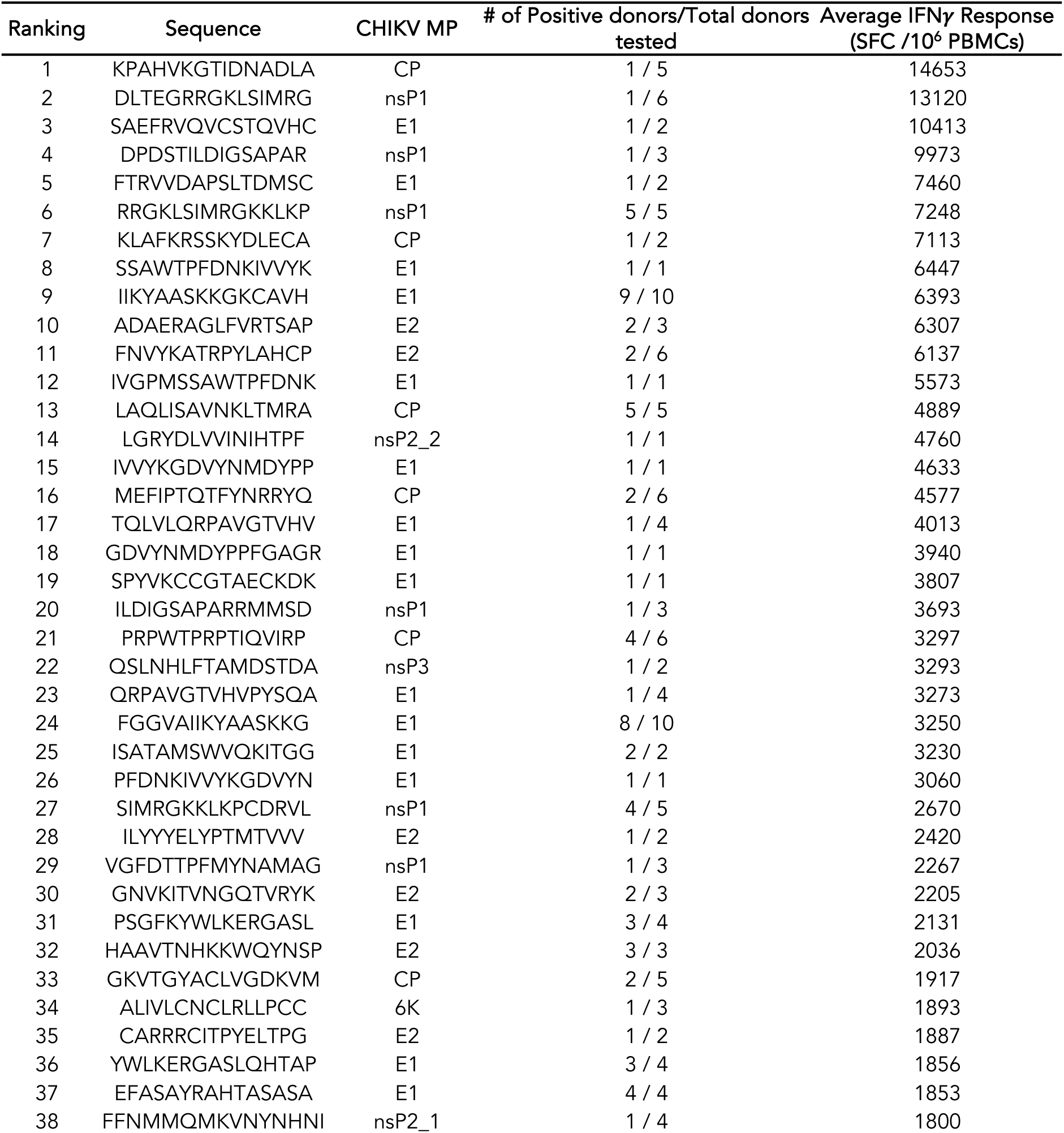
Epitopes that elicit 75% of total IFN*γ* response.

Specific epitopes were identified in 13/15 donors, with each donor recognizing on average 12 epitopes (range: 0-25 epitopes; Figure 3C). As expected in a heterogenous MHC donor cohort, 75% of epitopes were identified in only one donor (Figure 3D). Four peptides were most frequently recognized, with two contiguous peptides – E1_1121-1135_ (FGGVAIIKYAASKKG) and E1_1126-1140_ (IIKYAASKKGKCAVH) – being recognized by the eight and nine donors, respectively. Both E1_1121-1135_ and E1_1126-1140_ peptides were recognized by the same eight donors. The other two commonly recognized peptides – CP_41-55_ (LAQLISAVNKLTMRA) and nsP1_221-235_ (RRGKLSIMRGKKLKP) - were recognized by five donors each (Figure 3D). All four of the commonly recognized epitopes also were part of the top 38 epitopes that elicited a high magnitude of response (Table 2).

### Mechanism of immunodominance for the nsP1_226-235,_ E1_1126-1140_ and CP_46-50_ regions

The data presented above pin-points residues nsP1_226-235_, E1_1126-1140_ and CP_46-50_ as being associated with remarkable frequency and magnitude of responses. Additional experiment addressed the molecular mechanisms associated with such dominance, and whether these regions would likely contain a single epitope encoded in the overlap between the two consecutive 15mer peptides, or whether they would happen to contain multiple distinct epitopes.

Accordingly, we synthetized series of 15mer peptides overlapping by 14 residues and spanning residues 216-245 for nsP1, 1116-1145 for E1, and 36-65 for CP. These peptides were tested for immunogenicity with a T cell line derived by two-week *in vitro* restimulation of PBMCs from chronic and recovered donors (fine epitope mapping cohort from Table 1). Plotting responses from all donors clearly indicated that the peptides associated with optimal response all share the KLSIMRGKKL, IKYAASKKG and LISAVNKLTMR core regions for nsP1 (Figure 3E), E1 (Figure 3F) and CP (Figure 3G), respectively. In addition, T cells from recovered individuals recognized the same core sequences for nsP1 and E1 proteins, but at a considerably lower magnitude as compared to chronic donors.

Next, we used the IEDB NetMHCIIpan 4.1 EL method^29^ to perform HLA binding predictions for the reference set of 27 most common HLA class II alleles which has been shown to provide a 100% coverage of the general population when combined^30^. Predictions were performed using the 15mer nsP1_221-235_ (RRGKLSIMRGKKLKP), E1_1123-1136_ (GVAIIKYAASKKGKC) and CP_42-56_ (LAQLISAVNKLTMRAV) epitopes that encompassed the experimentally identified core motif, flanked by two and one additional amino acids at the N- and C-termini, respectively. The IEDB-recommended percentile value of less than 20% was used as the threshold defining a predicted binding event. As shown in Tables 3, 4 and 5, remarkably the nsP1 and E1 epitope was predicted to bind 18/27 (67%) and 19/27 (70%) of the HLA molecules, respectively, with frequency of predicted binding being slightly lower at 11 (44%) HLA alleles for the CP epitope. For each of the epitopes, we also tested the binding capacity of two negative control 15mers (nsP1: DLTEGRRGKLSIMRG and KKLKPCDRVLFSVGS; E1: THSSDFGGVAIIKYA and ASKKGKCAVHSMTNA; CP: RQAGQLAQLISAVNK and LTMRAVPQQKPRRNR) located at the N and C termini of the epitope region (nsP1: Table S2A and S2B; E1: Tables S3A and S3B; CP: Tables S4A and S4B). In the case of the control peptides, binding capacity was predicted for 0/27 (0%), and 3/27 (11%) HLA alleles for N- and C-termini nsP1 peptides respectively (Figure 3E), and 3/27 HLA (11%) and 0/27 (0%) HLA alleles for N- and C-terminus E1 peptides respectively (Figure 3F). For CP, 3/27 (11%) and 1/27(4%) HLA alleles were predicted to bind to N and C-terminus peptide (Figure 3G). A chi-square test was performed to examine if there was a statistical difference in binding for the finely mapped epitope and control peptides. For all proteins, a significant difference was found between predicted binding of peptides (nsP1, E1: p < 0.0001; CP: p = 0.0037). Additionally, we sequenced the HLA locus for each of the tested donors in our cohort using PCR amplification of genomic DNA and results are listed in Table S5. Each donor contained at least one HLA allelic variant that was predicted to be a strong binder for the peptides tested.

**Table 3:**
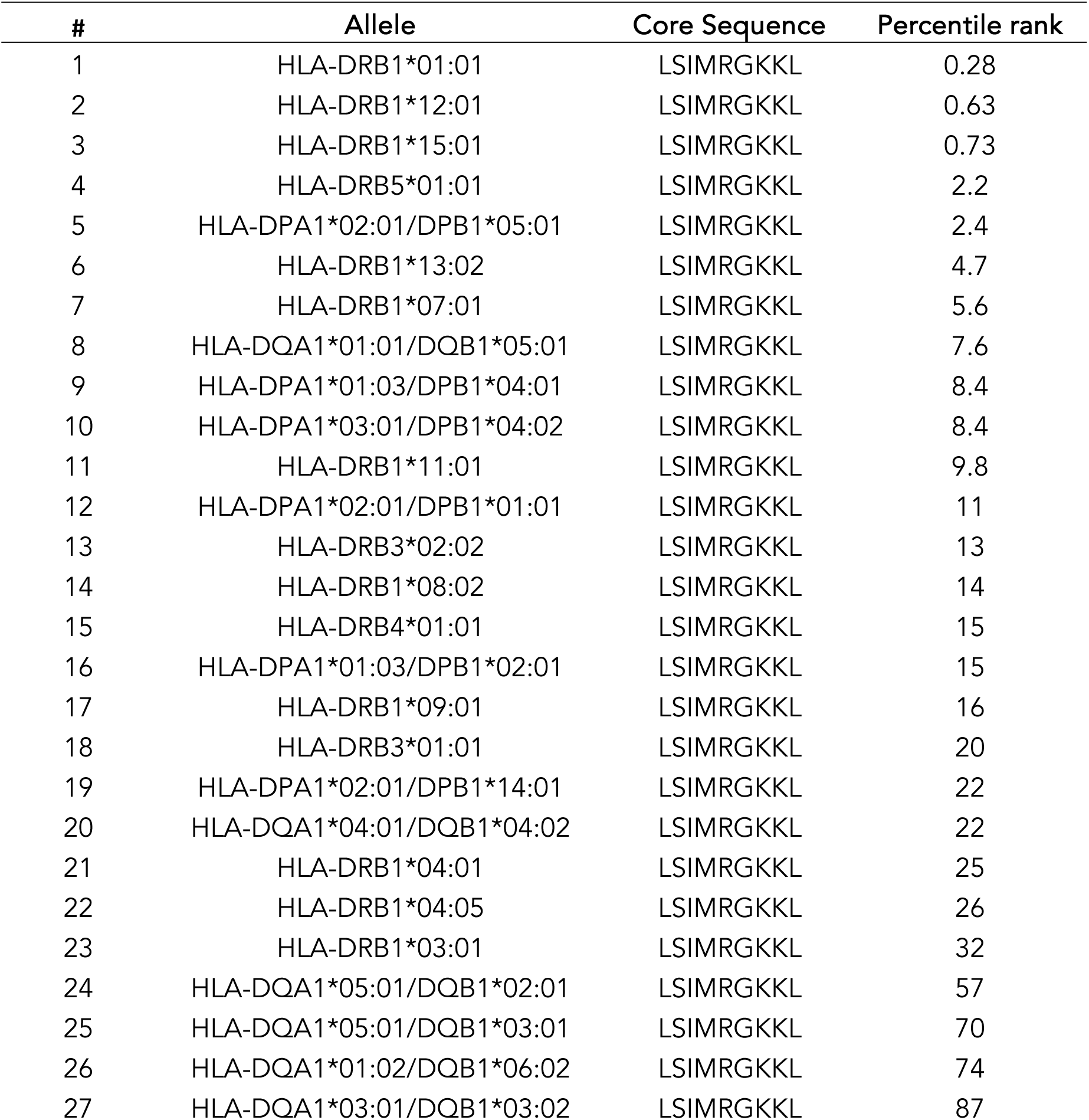
HLA binding predictions for the finely mapped epitope sequence in the nsP1 protein.

| # | Allele | Core Sequence | Percentile rank |
| --- | --- | --- | --- |
| 1 | HLA-DRB1*01:01 | LSIMRGKKL | 0.28 |
| 2 | HLA-DRB1*12:01 | LSIMRGKKL | 0.63 |
| 3 | HLA-DRB1*15:01 | LSIMRGKKL | 0.73 |
| 4 | HLA-DRB5*01:01 | LSIMRGKKL | 2.2 |
| 5 | HLA-DPA1*02:01/DPB1*05:01 | LSIMRGKKL | 2.4 |
| 6 | HLA-DRB1*13:02 | LSIMRGKKL | 4.7 |
| 7 | HLA-DRB1*07:01 | LSIMRGKKL | 5.6 |
| 8 | HLA-DQA1*01:01/DQB1*05:01 | LSIMRGKKL | 7.6 |
| 9 | HLA-DPA1*01:03/DPB1*04:01 | LSIMRGKKL | 8.4 |
| 10 | HLA-DPA1*03:01/DPB1*04:02 | LSIMRGKKL | 8.4 |
| 11 | HLA-DRB1*11:01 | LSIMRGKKL | 9.8 |
| 12 | HLA-DPA1*02:01/DPB1*01:01 | LSIMRGKKL | 11 |
| 13 | HLA-DRB3*02:02 | LSIMRGKKL | 13 |
| 14 | HLA-DRB1*08:02 | LSIMRGKKL | 14 |
| 15 | HLA-DRB4*01:01 | LSIMRGKKL | 15 |
| 16 | HLA-DPA1*01:03/DPB1*02:01 | LSIMRGKKL | 15 |
| 17 | HLA-DRB1*09:01 | LSIMRGKKL | 16 |
| 18 | HLA-DRB3*01:01 | LSIMRGKKL | 20 |
| 19 | HLA-DPA1*02:01/DPB1*14:01 | LSIMRGKKL | 22 |
| 20 | HLA-DQA1*04:01/DQB1*04:02 | LSIMRGKKL | 22 |
| 21 | HLA-DRB1*04:01 | LSIMRGKKL | 25 |
| 22 | HLA-DRB1*04:05 | LSIMRGKKL | 26 |
| 23 | HLA-DRB1*03:01 | LSIMRGKKL | 32 |
| 24 | HLA-DQA1*05:01/DQB1*02:01 | LSIMRGKKL | 57 |
| 25 | HLA-DQA1*05:01/DQB1*03:01 | LSIMRGKKL | 70 |
| 26 | HLA-DQA1*01:02/DQB1*06:02 | LSIMRGKKL | 74 |
| 27 | HLA-DQA1*03:01/DQB1*03:02 | LSIMRGKKL | 87 |

**Table 4:** HLA binding predictions for the finely mapped epitope sequence in the E1 protein.

| # | Allele | Core Sequence | Percentile Rank |
| --- | --- | --- | --- |
| 1 | HLA-DRB1*15:01 | IIKYAASKK | 1.1 |
| 2 | HLA-DRB5*01:01 | IIKYAASKK | 1.2 |
| 3 | HLA-DRB1*09:01 | IKYAASKKG | 4.2 |
| 4 | HLA-DRB1*08:02 | IKYAASKKG | 4.4 |
| 5 | HLA-DQA1*04:01/DQB1*04:02 | IKYAASKKG | 4.8 |
| 6 | HLA-DPA1*02:01/DPB1*05:01 | IIKYAASKK | 4.9 |
| 7 | HLA-DRB1*11:01 | IKYAASKKG | 6 |
| 8 | HLA-DRB1*13:02 | IIKYAASKK | 6.8 |
| 9 | HLA-DRB1*04:01 | IKYAASKKG | 7 |
| 10 | HLA-DRB1*07:01 | IKYAASKKG | 7.5 |
| 11 | HLA-DRB1*01:01 | IKYAASKKG | 8.7 |
| 12 | HLA-DQA1*01:02/DQB1*06:02 | IKYAASKKG | 9.5 |
| 13 | HLA-DQA1*05:01/DQB1*03:01 | IKYAASKKG | 9.7 |
| 14 | HLA-DRB1*12:01 | IIKYAASKK | 13 |
| 15 | HLA-DRB4*01:01 | IKYAASKKG | 14 |
| 16 | HLA-DPA1*01:03/DPB1*02:01 | VAIIKYAAS | 14 |
| 17 | HLA-DRB3*02:02 | IKYAASKKG | 17 |
| 18 | HLA-DRB1*04:05 | IKYAASKKG | 17 |
| 19 | HLA-DPA1*02:01/DPB1*14:01 | IKYAASKKG | 20 |
| 20 | HLA-DQA1*01:01/DQB1*05:01 | IIKYAASKK | 21 |
| 21 | HLA-DRB1*03:01 | IIKYAASKK | 24 |
| 22 | HLA-DPA1*03:01/DPB1*04:02 | VAIIKYAAS | 27 |
| 23 | HLA-DPA1*01:03/DPB1*04:01 | VAIIKYAAS | 28 |
| 24 | HLA-DPA1*02:01/DPB1*01:01 | IIKYAASKK | 29 |
| 25 | HLA-DRB3*01:01 | IIKYAASKK | 49 |
| 26 | HLA-DQA1*05:01/DQB1*02:01 | IKYAASKKG | 50 |
| 27 | HLA-DQA1*03:01/DQB1*03:02 | IKYAASKKG | 68 |

**Table 5:** HLA binding predictions for the finely mapped epitope sequence in the CP protein.

| # | Allele | Core Sequence | Percentile Rank |
| --- | --- | --- | --- |
| 1 | HLA-DRB1*11:01 | ISAVNKLTM | 2.7 |
| 2 | HLA-DRB1*12:01 | ISAVNKLTM | 2.9 |
| 3 | HLA-DRB1*13:02 | ISAVNKLTM | 3.5 |
| 4 | HLA-DRB1*08:02 | ISAVNKLTM | 4.5 |
| 5 | HLA-DPA1*02:01/DPB1*05:01 | ISAVNKLTM | 4.8 |
| 6 | HLA-DRB1*03:01 | ISAVNKLTM | 5.3 |
| 7 | HLA-DRB1*15:01 | ISAVNKLTM | 13 |
| 8 | HLA-DPA1*02:01/DPB1*01:01 | ISAVNKLTM | 15 |
| 9 | HLA-DPA1*01:03/DPB1*04:01 | ISAVNKLTM | 17 |
| 10 | HLA-DPA1*03:01/DPB1*04:02 | ISAVNKLTM | 19 |
| 11 | HLA-DPA1*01:03/DPB1*02:01 | ISAVNKLTM | 20 |
| 12 | HLA-DRB5*01:01 | ISAVNKLTM | 20 |
| 13 | HLA-DRB4*01:01 | ISAVNKLTM | 22 |
| 14 | HLA-DRB1*01:01 | LISAVNKLT | 22 |
| 15 | HLA-DRB1*09:01 | LISAVNKLT | 23 |
| 16 | HLA-DRB3*01:01 | ISAVNKLTM | 27 |
| 17 | HLA-DRB1*04:01 | LISAVNKLT | 28 |
| 18 | HLA-DRB3*02:02 | LISAVNKLT | 29 |
| 19 | HLA-DRB1*07:01 | ISAVNKLTM | 31 |
| 20 | HLA-DPA1*02:01/DPB1*14:01 | ISAVNKLTM | 41 |
| 21 | HLA-DQA1*04:01/DQB1*04:02 | ISAVNKLTM | 42 |
| 22 | HLA-DRB1*04:05 | LISAVNKLT | 43 |
| 23 | HLA-DQA1*05:01/DQB1*03:01 | LISAVNKLT | 44 |
| 24 | HLA-DQA1*01:02/DQB1*06:02 | LISAVNKLT | 49 |
| 25 | HLA-DQA1*01:01/DQB1*05:01 | ISAVNKLTM | 50 |
| 26 | HLA-DQA1*05:01/DQB1*02:01 | ISAVNKLTM | 53 |
| 27 | HLA-DQA1*03:01/DQB1*03:02 | ISAVNKLTM | 61 |

Altogether, these data suggest that the nsP1_226-235,_ E1_1126-1140_, and CP_46-50_ regions contain a single epitope, characterized by KLSIMRGKKL, IKYAASKKG and LISAVNKLTMR core regions, respectively. The regions are associated with a remarkable capacity to bind many or most human HLA class II molecules frequently expressed in the worldwide population, thus explaining their dominance.

### Development and validation of an epitope-based CHIKV CD4^+^ T cell MP

The MPs utilized so far were based on pools of overlapping peptides and variants spanning the entire sequence of the CHIKV proteome. Having now identified actual epitopes recognized by human CD4^+^ T cells, we leveraged this knowledge to create a CHIKV-specific MP containing the verified epitopes that thus encompassed fewer peptides. Accordingly, we designed two MPs - structural and non-structural - that consisted of 123 consensus epitopes and 24 variant previously identified epitopes. The structural epitope MP, referred to as CHIKV S, and the non-structural epitope MP, referred to as CHIKV NS, consisted of 84 and 63 epitopes, respectively. We envision that such epitope-based MP, by analogy with those developed for other indications such as DENV^31^, pertussis^32^, tuberculosis^33^ or SARS-CoV-2^34^ could be used to study CD4^+^ T cell responses if only small amounts of blood are available, as typical in pediatric and clinical studies.

To validate these epitope-based CHIKV MPs, we recruited an independent cohort of 19 CHIKV-seropositive donors with chronic CHIKVD (referred to as “chronic”) and seven CHIKV-seronegative or uninfected controls. Cohort characteristics are provided in Table 1. Infections were confirmed by measuring CHIKV-specific IgG titers (Figure S1A). We stimulated PBMCs with CHIKV S, CHIKV NS and a combined CHIKV S+NS MPs for 24h and measured antigen-specific CD4^+^ T cell responses via upregulation of OX40+CD137+. CHIKV-specific CD4^+^ T cell responses were not detected in uninfected controls; however, 8/19 (42%) and 5/19 (26%) of chronic CHIKVD donors were positive in responses to individual CHIKV S and CHIKV NS MPs, respectively (Figure 4A). Following stimulation with combined CHIKV S+NS MP (147 peptides), we detected a higher number of CHIKV-specific CD4^+^ T cell positive responses in 11/19 (58%) of donors. In comparison, 12/17 (65%) donors were found to be positive in the initial screening upon averaging responses to all the overlapping peptide MPs that spanned the entire CHIKV proteome (992 peptides). The 7-fold reduction in the number of peptides required for screening and therefore the lower amount of blood volumes necessary points to an advantage in the feasibility of identifying the CHIKV-specific T cell responses in low amounts of blood, typically available in clinical and vaccine cohorts.

**Figure 4:**
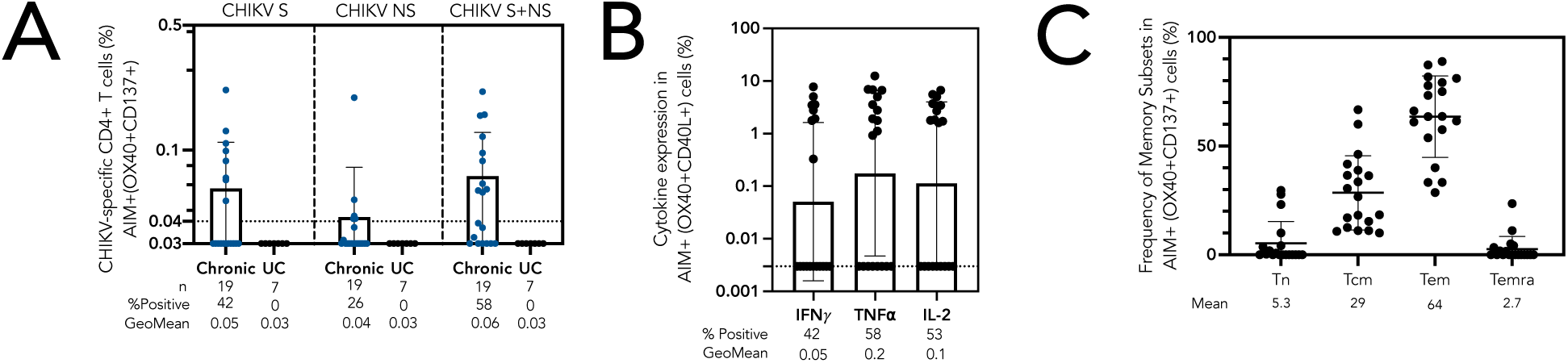
CHIKV CD4^+^ T cell specific epitope megapool induces a robust response. **(A)** Frequency of antigen-specific CD4^+^ T cells quantified by the AIM assay (OX40^+^CD137^+^) after 24-hour stimulation with CHIKV epitope megapools consisting of epitopes from structural (S), non-structural (NS) and combined structural and non-structural (CHIKV S+NS) proteins in 19 chronic CHIKV donors (chronic; blue dots) and seven CHIKV seronegative or uninfected donors (black dots; uninfected controls (UC)), with n representing the number of donors. % Positive refers to the percent of donors who are above the LOS (0.04), indicated by the dotted line. **(B)** Frequency of specific cytokine producing cells (IFN*γ*, TNF⍺ and IL-2) from the CHIKV-specific CD4^+^ T cells (AIM+ OX40^+^CD40L^+^) after stimulation with the combined structural and non-structural CHIKV epitope megapool (CHIKV S+NS) in 19 chronic CHIKV donors. % Positive refers to the percent of donors that are above the LOS (0.003), indicated by the dotted line. **(C)** Frequency of CHIKV-specific AIM+ CD4^+^ T cell memory subsets (AIM+ OX40^+^CD137^+^) in 19 chronic CHIKV donors post-stimulation with the CHIKV S+NS epitope MP, based on the expression of CCR7 and CD45RA in AIM+ CD4^+^ T cells as: T naive (CCR7^+^ CD45RA^+^), TCM (T central memory; CCR7^+^ CD45RA^-^), TEM (T effector memory; CCR7^-^ CD45RA^-^) and TEMRA (T effector memory re-expressing CD45RA; CCR7^-^ CD45RA^+^) Data are represented as mean ± SD or geomean ± geometric SD.

Next, within the AIM+ specific responses, we measured cytokine expression of IFN*γ*, tumor necrosis factor-alpha (TNF⍺) and interleukin 2 (IL-2) after simulation with the combined CHIKV S+NS MP in chronic CHIKV-seropositive donors via the intracellular staining assay. 58% and 53% of AIM+ (OX40+CD40L+) donors were positive for TNF⍺ and IL-2 expression, respectively, while 42% of donors were positive for IFN*γ* expression (Figure 4B). The majority of CHIKV S+NS-specific AIM+ CD4^+^ T cells displayed T effector memory phenotype (TEM: CD45RA-CCR7+), followed by T central memory cell phenotype (TCM: CD45RA-CCR7-) (Figure 4C).

### Sequence conservation of CHIKV CD4^+^ T cell epitopes in arthritogenic and encephalitic alphaviruses

Next, we investigated to what degree CHIKV epitopes identified in this study were conserved across other arthritogenic and encephalitic alphaviruses. We selected a representative set of viruses from the alphavirus genus (Figure 5A) and retrieved the associated protein sequences from Virus Pathogen Resource database (ViPR; www.viprbrc.org)^35^. We then calculated the percentage of conservation of each CHIKV peptide using the Conservation Analysis tool in IEDB-AR^36^. Plotting the median percent conservation for arthritogenic (excluding CHIKV sequences; n = 8) and encephalitic alphaviruses (n = 8) for all CHIKV peptides studied herein revealed regions of high conservation in the CP, nsP4, nsP1 and nsP2 proteins and regions of low conservation in the 6K, E2 and E1 proteins in both arthritogenic and encephalitic groups (Figure 5B). As expected on the basis of closer phylogenetic relationships, the degree of conservation of CHIKV peptides was higher for all proteins in arthritogenic alphaviruses compared to encephalitic alphaviruses. Interestingly, the nsP4 protein showed the second highest degree of conservation in both arthritogenic and encephalic groups, however, few epitopes were experimentally identified in that region.

**Figure 5:**
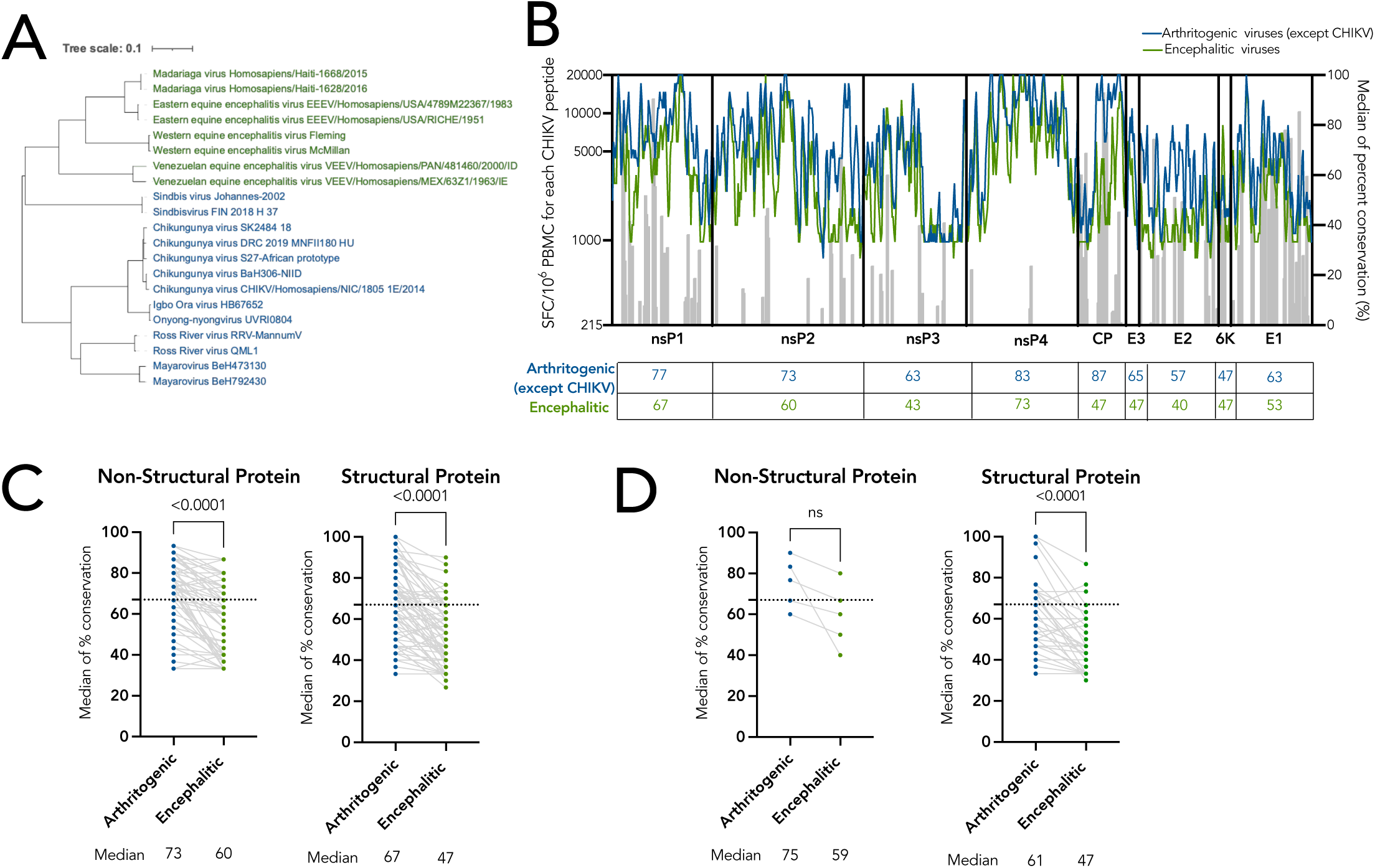
Sequence conservation of CHIKV proteome in arthritogenic and encephalitic alphaviruses. **(A)** Phylogenetic tree indicating the viral sequences used to calculate the percentage of conservation of CHIKV peptides. The tree is divided into arthritogenic (blue) and encephalitic (green) alphaviral sub-groups. **(B)** Magnitude of response and median percent conservation of CHIKV peptides in arthritogenic (blue) and encephalitic (green) alphaviruses. The left y-axis and gray bars refer to the magnitude of response of each CHIKV epitope. The right y-axis, and blue and green lines refer to median percent conservation for each CHIKV peptide. The solid lines separate each CHIKV protein. The median of percent conservation for each CHIKV protein is shown in the table below for arthritogenic (except CHIKV) and encephalitic alphaviruses. **(C)** Median percent conservation of all CHIKV epitopes in arthritogenic (except CHIKV sequences; blue) and encephalitic (green) alphaviruses separated based on non-structural (nsP1, nsP2, nsP3 and nsP4) and structural (CP, E3, E2, 6K and E1) proteins. Each dot indicates a CHIKV epitope. The median percent conservation for each sub-group is shown below. **(D)** Median percent conservation of each immunodominant epitope (recognized by two or more donors) in arthritogenic (except CHIKV sequences; blue) and encephalitic (green) alphaviruses separated based on non-structural (nsP1, nsP2, nsP3 and nsP4) and structural (CP, E3, E2, 6K and E1) proteins. For (C) and (D), the dotted lines refer to 67% threshold that has previously been shown to define cross-reactive epitopes. The median percent conservation for each sub-group is shown below. Data were analyzed for statistical significance using Wilcoxon signed-rank test (p>0.05, ns = nonsignificant).

For the 123 individual consensus CHIKV epitopes and 24 variant epitopes identified in this study, we plotted the median percent conservation in arthritogenic and encephalitic alphaviruses (Figure 5C). Overall, the median percent conservation for both non-structural and structural epitopes was significantly higher in the arthritogenic group (non-structural: 73%, structural: 67%) compared to in the encephalitic group (non-structural: 60%; structural: 47%) (p < 0.0001).

Previous studies have experimentally defined 67% conservation to be associated with cross-reactivity between SARS-CoV-2 epitopes and common-cold coronaviruses^25,26^. As such, we used 67% median conservation as the threshold to identify CHIKV CD4^+^ T cell epitopes that could potentially be cross-reactive with other alphaviruses. Overall, 68% (42/62) of all CHIKV non-structural and 47% (35/75) of structural epitopes were predicted to be potentially cross-reactive with other arthritogenic viruses and 31% (19/62) of the CHIKV non-structural and 13% (10/75) of the structural epitopes were predicted to be potentially cross-reactive with other encephalitic viruses (Figure 5C).

We further analyzed the five epitopes in the non-structural proteins and 35 epitopes in the structural proteins that were recognized in two or more donors (referred to as immunodominant) (Figure 5D). Four out of five dominant epitopes (80%) in the non-structural proteins were predicted to be potentially cross-reactive with other viruses in arthritogenic versus one out of four (25%) in the encephalitic group. Out of the 35 immunodominant epitopes in the structural proteins, 13/35 (37%) were predicted to be potentially cross-reactive with other viruses in arthritogenic versus 3/25 (12%) in the encephalitic groups, with overall median conservation of all immunodominant epitopes significantly higher in the arthritogenic than in encephalitic groups (p < 0.0001). The potential cross-reactivity across other representatives of the alphavirus genus could be important in the context of CHIKV infection or vaccination.

## DISCUSSION

Here, we investigated the repertoire of CHIKV-specific CD4^+^ T cell responses in individuals with chronic CHIKVD. To the best of our knowledge, this is the first study to report human CHIKV-specific CD4^+^ T cells epitopes. Epitopes in E1 and nsP1 proteins accounted for nearly half of the 123 identified epitopes and more than half of the total magnitude of the response. Specifically, two regions in nsP1_226-235_ and E1_1126-1140_ proteins were identified as immunodominant and were marked by high magnitude of response. Fine epitope mapping determined that these regions each contained a single epitope recognized in the same frame by multiple donors. The immunodominance of these two regions could be explained by the observation that these core regions are predicted to bind multiple HLA allelic variants that are frequently expressed in the world population. Beyond E1 and nsP1, CP contained the next highest number of epitopes, which accounted for 18% of the overall response. Interestingly, CP also contained a single epitope with the ability to bind to multiple HLA alleles. Thus, overall, nsP1, E1 and CP proteins were the most immunogenic, highlighting their importance for vaccine development.

While no CHIKV-specific T cell epitopes had been previously identified in humans, a few studies have analyzed CHIKV-specific T cell targets in mice. Interestingly, nine of the 26 epitopes defined in CHIKV-infected C57BL/6 mice overlap with the epitopes identified in our study, which is remarkable given the highly distinct MHC in humans and mice^20^. Specific sequences of the two pathogenic CD4^+^ T cell targets discovered in mice in a separate study by Teo et al. were found to be immunogenic in our study as well^18^. This suggests that murine models can be a representative model to study CHIKV T cell responses relevant for human disease.

We also assessed conservation of the T cell targets of CHIKV across other alphaviruses, which is relevant in the context of potential universal alphaviral vaccines^37^. Our analysis revealed regions of broad conservation of the CHIKV proteome in arthritogenic and encephalitic alphaviruses. While the nsP4 protein had 84% median conservation in arthritogenic alphaviruses, only three epitopes were identified in the nsP4 antigen. On the other hand, structural proteins, specifically CP, E2 and E1 proteins, which elicited a high magnitude of response with a high number of immunogenic epitopes, were associated with lower degree of conservation, indicating lower potential cross-reactivity with other alphaviruses. Importantly, however, highest degree of conservation was found in immunodominant epitopes in non-structural proteins of the arthritogenic and encephalitic alphaviruses. Based on these results, the design of a universal alphavirus vaccine could rely on focusing T cell responses on the relatively few epitopes of the viral proteome that are highly conserved and also immunogenic for human T cell responses.

Overall, our work provides an in-depth characterization of CHIKV CD4^+^ T cell epitopes in chronic CHIKVD and extends the analysis to determine conservation of the identified epitopes in arthritogenic and encephalitic alphaviruses. Additionally, we designed a CHIKV-specific CD4 epitope MP which will aid in the characterization of CD4^+^ T cell response with small blood volumes. Our study has broadened our understanding of CHIKV-specific T cell responses, which will enable monitoring of human CD4+ T cell responses in the context of study of CHIKV immunopathology and in the context of alphavirus vaccine development.

### Limitations of the study

There are several limitations that need to be considered when interpreting our findings. Due to limited availability of samples, we were unable to study CD4^+^ T cell targets during acute phase of CHIKV infection. Our studies are limited in terms of number of individuals studied, and all were derived from a single geographical region. Future work could focus on a more exhaustive epitope screening in additional individuals and cohorts from diverse geographical origins. Furthermore, we defined potentially cross-reactive epitopes in alphaviruses based on purely *in-silico* analysis of CHIKV epitopes. To truly determine the regions of CHIKV that could be cross-reactive with other alphaviruses, experimental testing of epitopes would be required.

## METHODS

### Study donor cohort

We enrolled 47 donors from Departamento del Atlántico on the Caribbean region of Colombia who had been diagnosed with CHIKV during the 2014-2015 epidemic. The criteria of a positive CHIKV diagnosis included: 1) whether the symptoms matched with CHIKV, 2) whether the patient lived in or visited a region where CHIKV had been detected using a RT-PCR assay, 3) whether other members of the patient’s household or other residents in the neighborhood had been diagnosed with CHIKV, 4) whether the infections with dengue virus, which often co-circulates with CHIKV and has similar clinical presentation, has been ruled out through laboratory tests such as ELISA and rapid diagnostic assays. Not all patient samples were subjected to laboratory tests to confirm CHIKV infection. Local medical professionals determined if the epidemiological link based on the criteria mentioned above was considered sufficient to diagnose a case as CHIKV positive. For the purpose of our studies, we confirmed infection by measuring the levels of anti-CHIKV IgG antibodies using the serum of all individuals (Figure S1A). 40 individuals were confirmed to be seropositive for CHIKV infection, while seven individuals were seronegative. Of the 40 seropositive individuals, 36 displayed chronic arthritis-like symptoms, such as joint pain and swelling, and the remaining four had recovered post-acute infection.

The cohort was divided into three groups – CHIKV epitope screening group, CHIKV fine epitope mapping group and CHIKV epitope megapool (MP) validation group. PBMCs from 17 CHIKV seropositive individuals with chronic CHIKVD were used to screen for epitopes, 10 of which were also used for fine epitope mapping. Responses to core regions for chronic donors were compared to responses from four recovered donors. PBMCs from the remaining 19 CHIKV seropositive individuals with chronic CHIKVD were used to validate the identified epitopes. Seven CHIKV seronegative donors were used as uninfected controls for epitope MP validation.

In addition, to determine the threshold of *in vitro* positive responses to CHIKV MPs, six uninfected donors were recruited from the National Blood Center of Nicaragua Red Cross.

At the time of enrollment in the study, all individual donors provided informed consent that their samples could be used for any future studies, including this study. All blood and serum samples were anonymized and given a code number. In addition to collecting blood and serum samples, information regarding the date of their CHIKV diagnosis, the date of symptom onset, the date of sample collection, their age, their gender, and the specific symptoms at the time of CHIKV diagnosis and sample collection was also collected for each participant. Information regarding sex, age, current symptoms and CHIKV serostatus for each cohort is provided in Table 1.

### Study approval

All samples were collected under an approved IRB protocol of La Jolla Institute for Immunology (LJI VD154) and Universidad del Norte, Colombia (Protocol number 153).

### PBMC isolation

Whole blood samples were collected from CHIKV donors in Colombia. Large volumes of the donors’ blood were collected in sterile bags which contained 3.27 g citric acid, 26.3 g NaH_2_-citrate and 2.22 g NaHCO3 per liter (Fresenius Kabi, Fresenius HemoCare, Brasil Ltd, Brasil) and then diluted by 50% v/v with RPMI-1640 media (31800-105, Gibco, USA) containing 0.2% w/v NaHCO_3_. 34 mL volumes were gently layered into multiple sterile 50 mL tubes (430290, Corning, USA) containing 12 mL of sterile endotoxin-tested Ficoll-Paque PLUS (1.077 g/ml density) (GE Healthcare Biosciences, Sweden). After centrifugation at 500xg for 30 min, the buffy-coat peripheral blood mononuclear cells (PBMCs) were collected from immediately above the Ficoll-Paque-plasma interface, diluted with 50% v/v RPMI 1640 medium and again centrifuged at 500xg to pellet the PBMCs. These PBMCs were then re-suspended in large volumes of RPMI 1640 medium containing 5% v/v fetal calf serum (FCS) (F-0926, Sigma, USA) and centrifuged again at 500xg for 30 min. After a further wash, PMBCs were re-suspended in ice-cold 10% dimethyl sulfoxide (D-2650, Sigma, USA) in FCS at approximately 20 x 10^6^ PBMCs/mL and 1 mL aliquots contained in 1.5 mL polypropylene cryo-vials (5000-1020: Nalgene System 100, Thermo Scientific, USA) were slowly (-1°C/min) frozen down to -80°C in a isopropanol alcohol-walled freezing unit (Nalgene Mr. Frosty: C-1562 Sigma-Aldrich, USA) before being transferred to 25-vial boxes in liquid nitrogen. Whole blood samples from CHIKV uninfected donors in Nicaragua were collected and isolated using the previously described methods^38^.

### Peptide pools

We retrieved 257 and 350 full-length CHIKV structural and non-structural polyprotein sequences from the Virus Pathogen Resource database (ViPR; www.viprbrc.org)^35^ using the following query: Chikungunya virus, Gene product name: structural OR non-structural polyprotein, Remove duplicate sequences. Unresolved sequences were removed. The query was performed in November 2019. Based on the results, we generate two separate consensus alignments for structural and non-structural polyproteins.

The number of sequences available varied as a function of geographic locations. To ensure balanced representation, the number of isolates by geographical region was limited to a maximum of 10. In total, 158 structural and 61 non-structural sequences were selected. For each polyprotein, sequences were aligned using MUSCLE, and consensus sequences were BLASTed to identify a representative isolate (GenBank ID: AQX78118.1 and AQX78116.1), using the tools hosted in the ViPR website. To account for variants from the consensus sequences, we identified an additional 105 and 146 amino acid variants, which were found in frequencies above 10% from structural and non-structural consensus sequences, respectively.

Based on the consensus sequence, we synthesized 741 15-mer peptides overlapping by 10 residues and an additional 251 15mers corresponding to the variant sequences. These peptides were resuspended in dimethyl sulfoxide (DMSO) and divided into 11 megapools (MP). The number of peptides per MP are listed in Table S1. The nsP2 antigen was split into two pools (nsP2_1 and nsP2_2) of 117 and 81 peptides, respectively.

The MP approach has been used previously described^24^. Large numbers of different epitopes are solubilized, pooled, and re-lyophilized to avoid cell toxicity problems associated with high concentrations of DMSO typically encountered when pooling after a single solubilization step.

Individual MP were further subdivided into mesopools (MS) encompassing 9-11 peptides, each of which were deconvoluted to identify individual epitopes. These number of MS corresponding to each MP ranged from 7 to 16, depending on the MP size. Introducing the MS intermediate screening step allowed to decrease the number of tests necessary to identify specific epitopes.

For epitope-based MPs, consensus and variant epitopes were synthesized, resuspended in DMSO post lyophilization and divided into structural and non-structural MPs, which were used for validation of epitope MPs. Each pool was tested 6-19 times.

### Activation induced Marker assay and Intracellular staining assay

PBMCs were thawed in 10 mL of RPMI 1640 (Corning) supplemented with 5% human AB serum (GeminiBio), penicillin [100 IU/mL], streptomycin [100μg/mL] (GeminiBio), and 2 mM L-glutamine (Gibco), and in the presence of benzonase (20μL/10mL). PBMCs were then plated at 1 x 10^6^ cells per well in 96-well U bottom plates and stimulated separately with each CHIKV-specific MP [1μg/mL] or CHIKV structural and non-structural epitope MP. An equimolar amount of DMSO was used for negative control. Stimulation with phytohemagglutinin (PHA, Roche) [1μg/mL] was used for positive controls [1μg/mL]. For samples that were tested for epitope identification, cells were stained for detection of activation-induced markers after 24h of stimulation.

For IntraCellular Staining (ICS) assays, 20h post-stimulation, Golgi-Plug and Golgi-Stop were added to the culture, in addition to anti-CD69 and anti-CD137 Abs. Cells were then washed, incubated with BD human FC block, and stained with LIVE/DEAD marker in the dark for 15 min. After incubation, cells were washed, surface stained in the dark for 30 min at 4°C, and then fixed with 1 % of paraformaldehyde (Sigma-Aldrich, St. Louis, MO). Subsequently, cells were permeabilized and stained with specific antibodies in the dark for 30 min at RT. All samples were acquired on Cytek Aurora. Antibodies used in the initial AIM screening assay are listed in Table S6 and the antibodies used in the AIM/ICS validation assay are listed in Table S7. A representative gating strategy for the AIM assay is shown in supplemental Figure S1B and for the combined AIM/ICS assays in Figure S2.

Antigen-specific CD4^+^ T cells were measured as a percentage of AIM+ (OX40+CD137+ and OX40+CD40L+) CD4^+^ T cells. The stimulation index (SI) was calculated by dividing the percentage of stimulated samples by those of the DMSO control. The limit of detection (LOD) was calculated as the value corresponding to the two-fold geometric 95% confidence interval of all DMSO values. Limit of sensitivity (LOS) was calculated using the median 2-fold standard deviation of all DMSO values. Response < LOD (0.02%) and SI < 2, after background subtraction, was normalized to the LOD. For initial AIM screening assay, SI> 2 and response > LOS (0.02%) was considered a positive response for antigen-specific CD4^+^ T cells. For the combined AIM/ICS assays, SI > 2 and response > LOS (0.04%) was considered a positive response for antigen-specific CD4^+^ T cells.

### T cell lines

PBMCs from AIM+ donors were cultured at 37°C and 5% CO2 in RPMI 1640 supplemented with 5% human serum at a density of 2 × 10^6^ cells per well in 6-well plates. The cells were stimulated individually with each MP that the donor was AIM positive for. Interleukin 2 (IL-2; 10 U/ml) was added every 3 to 4 days until cell harvest at day 14 (for mesopool evaluation) or day 17 (for individual peptide evaluation).

For fine epitope mapping, the cells were individually stimulated with either nsP1_221-235_ or nsP1_226-240_ or E_1121-1135_ and E1_1126-1140_ or CP_41-55_ and CP_46-60_ peptides on day 1 and restimulated on day 15 with individual peptides that overlapped by 14 residues and covered the entire region flanking the immunodominant regions (216-245 residues for nsP1, 1116-1145 residues for E1, and 36-65 residues for CP).

### FluoroSpot assay

CHIKV-specific responses were assessed by gamma interferon (IFN*γ*) FluoroSpot assay, following *in vitro* expansion at day 14, to detect responses to MS or individual peptides overlapping by 14 residues. For epitope screening, at day 17, positive pools were deconvoluted by FluoroSpot using individual CHIKV peptides (10 μg/mL) contained in the positive pool. Briefly, polyvinylidene fluoride (PVDF) plates (Millipore) were coated with anti-human IFN*γ* (1-D1K; Mabtech), and cells were plated in triplicate at 1 × 10^5^ cells per well. Cells were stimulated with either peptide pools (1 μg/mL) or individual peptides (10 μg/mL) in 0.1 mL of complete RPMI and incubated for 24 h at 37°C and 5% CO_2_.

Following stimulation, cells were discarded, and plates were incubated with biotinylated IFN*γ* monoclonal antibodies (MAb) (7-B6-1; Mabtech) for 2 h at 37°C and developed as described previously^39^. Positive responses were identified as those having >215 spot-forming cells (SFCs) per million cells, a SI of >2, and a p-value < 0.05 when compared to unstimulated cells using a t-test as previously described^40^.

### Serologic assays

Site-specifically biotinylated CHIKV E1 DIII and Halo-tag control antigens were coupled to unique MagPlex®-Avidin microspheres at a concentration of 5μg of antigen per 10^6^ beads in assay buffer (1% BSA + phosphate-buffered saline (PBS), pH 7.4) for 1h at 37°C with shaking at 700 rpm as described before^41^. Antigen coupled beads were washed and aliquoted 2,500 beads per antigen per well into a 96-well assay plate. Heat-inactivated (56°C for 30 min) human serum samples diluted at 1:500 in assay buffer were incubated with beads for 1h at 37°C with shaking at 700 rpm. After washing the beads with assay buffer, PE conjugated goat anti-human IgG Fc secondary Ab was added at 6μg/mL (Southern Biotech, catalog: 2014-09) and incubated for 1h at 37°C with shaking at 700 rpm. Beads were washed and resuspended in 100μL assay buffer for fluorescence analysis using the Luminex 200 system.

### HLA typing

HLA typing of Class I and Class II alleles of PBMCs of all donors were performed using locus-specific PCR amplification of genomic DNA by a laboratory accredited by the American Society for Histocompatibility and Immunogenetics at Murdoch University (Western Australia) as previously described^42^.

### Analysis of conservation of epitopes and non-epitopes in alphaviruses

Sequences of all alphaviruses were extracted from ViPR (www.viprbrc.org)^35^ based on sequence names provided in the phylogenetic tree in Figure 4A. Sequences were aligned using the online MAFFT version 7 alignment tool (https://mafft.cbrc.jp/alignment/server/index.html) and Neighbor-joining method (with default settings) was used to create a tree. The tree was visualized using Interactive Tool of Life (ITOL) web server. To calculate conservation, viruses were divided into arthritogenic, and encephalitic subgroups based on their main pathogenic characteristics. IEDB epitope conservancy tool^43^, with the threshold of 10%, was used to determine percent identity of each CHIKV peptide in the extracted alphaviral sequences. Median of percent conservation was calculated separately for encephalitic and arthritogenic alphaviruses.

## QUANTIFICATION AND STATISTICAL ANALYSIS

Data and statistical analyses were performed using FlowJo 10.8.1 and GraphPad Prism 9 (La Jolla, CA). The statistical details of the experiments are provided in the respective figure legends and in method details.

## Acknowledgements

We thank all members of the Weiskopf and Sette labs for their helpful discussions and all the subjects in Colombia and Nicaragua for participating in the study. **Funding**: This work was funded by National Institute of Health (NIH) Contract No. 75N93019C00065 to A.S. and D.W. and NIH Contract Number: 75N93024C00056 to A.S. and A.G.

## Author contributions

Conceptualization: D.W.; Methodology: R.A., C.H. and D.W.; Formal analysis: R.A., C.H. and D.W.; Investigation: R.A., C.H., F.H.C., Y.L., A.M.P., R.I.G., I.N.C., L.P., A.G.; Resources: E.H., A.B., A.G., C.M.R., A.K.F.; Writing – Original Draft: R.A. and D.W.; Writing – Review & Editing: R.A., C.H., F.H.C., Y.L., A.M.P., R.I.G., I.N.C., A.B., E.H., C.M.R., L.P., A.K.F., A.G., A.S. and D.W; Visualization: R.A., C.H., and D.W.; Funding acquisition: A.S. and D.W.; Supervision: D.W.

## Materials and Correspondence

Upon specific request and execution of material transfer agreement (MTA) to the lead contact, aliquots of the peptide pools used in this study will be made available. There are restrictions to the availability of the peptide reagents due to cost and limited quantity. All correspondence should be addressed to the lead contact.

## Data availability

All data reported in this paper will be shared by the lead contact upon request. The identified epitope sequences have been submitted to IEDB (submission IDs 1000910, 1000911 and 1000913).

## SUPPLEMENTAL FIGURE LEGENDS

**Figure S1: Cohort serology, representative gating strategy, AIM+ CD4^+^ T cell responses and FluoroSpot Assay threshold of positivity**

**(A)** Net CHIKV-specific IgG titers measured in the study cohort. Left panel shows titers for samples collected from Nicaragua (threshold - 10). Right panel shows titers for samples collected from Colombia (threshold – 907). The threshold (dotted line) was used to determine seropositivity and to confirm CHIKV infection. **(B)** Representative gating strategy to define antigen-specific CD3+CD4^+^ cells by the AIM assay. The cells shown here were stimulated with nsP1 protein. **(C)** Antigen-specific CD4+ T cells quantified by AIM (left: OX40+CD137+ and right: OX40+CD40L+) after 24-hour stimulation with all CHIKV MPs in 17 CHIKV seropositive donors with chronic CHIKVD. The dotted line represents the limit of sensitivity (LOS; 0.02%). Data are represented as geomean ± geometric SD. **(D)** Graph shows the IFN*γ* producing SFC/10^6^ PBMCs for six CHIKV-seronegative donors post *in vitro* stimulation with all CHIKV mesopools. Responses were plotted in descending order of magnitude of response and the top 1% response (red line) was calculated. 99% of the response was lower than 215 SFC/10^6^ PBMCs (dotted line), which was used as a threshold of positivity for following experiments.

**Figure S2: Representative gating strategy for the AIM/ICS assay**

Representative gating strategy to define antigen-specific CD4+ T cells (OX40+CD137 and Ox40+CD40L+) and their functional profile via ICS staining of cytokines (IFN*γ*, TNF⍺ and IL-2). Cells shown here were stimulated with the combined CHIKV S+NS epitope MP for initial gating and AIM gating, except with PHA for the cytokines.

**Table S1: Variant epitopes identified**

**Table S2A: HLA binding predictions for the N-term peptide in the immunodominant nsP1 protein region**

**Table S2B: HLA binding predictions for the C-term peptide in the immunodominant nsP1 protein region**

**Table S3A: HLA binding predictions for the N-term peptide in the immunodominant E1 protein region**

**Table S3B: HLA binding predictions for the C-term peptide in the immunodominant E1 protein region**

**Table S4A: HLA binding predictions for the N-term peptide in the immunodominant CP protein region**

**Table S4B: HLA binding predictions for the C-term peptide in the immunodominant CP protein region**

**Table S5: HLA alleles of the donors from the fine mapping epitope cohort Table S6: Antibodies used in the AIM screening assay**

**Table S7: Antibodies used in the AIM/ICS validation assay**

