## Supplemental Material for "Identification of immunogenic and cross-reactive chikungunya virus-specific CD4^+^ T cell epitopes in chronic chikungunya viral arthritic disease in humans"

### FIGURE S1

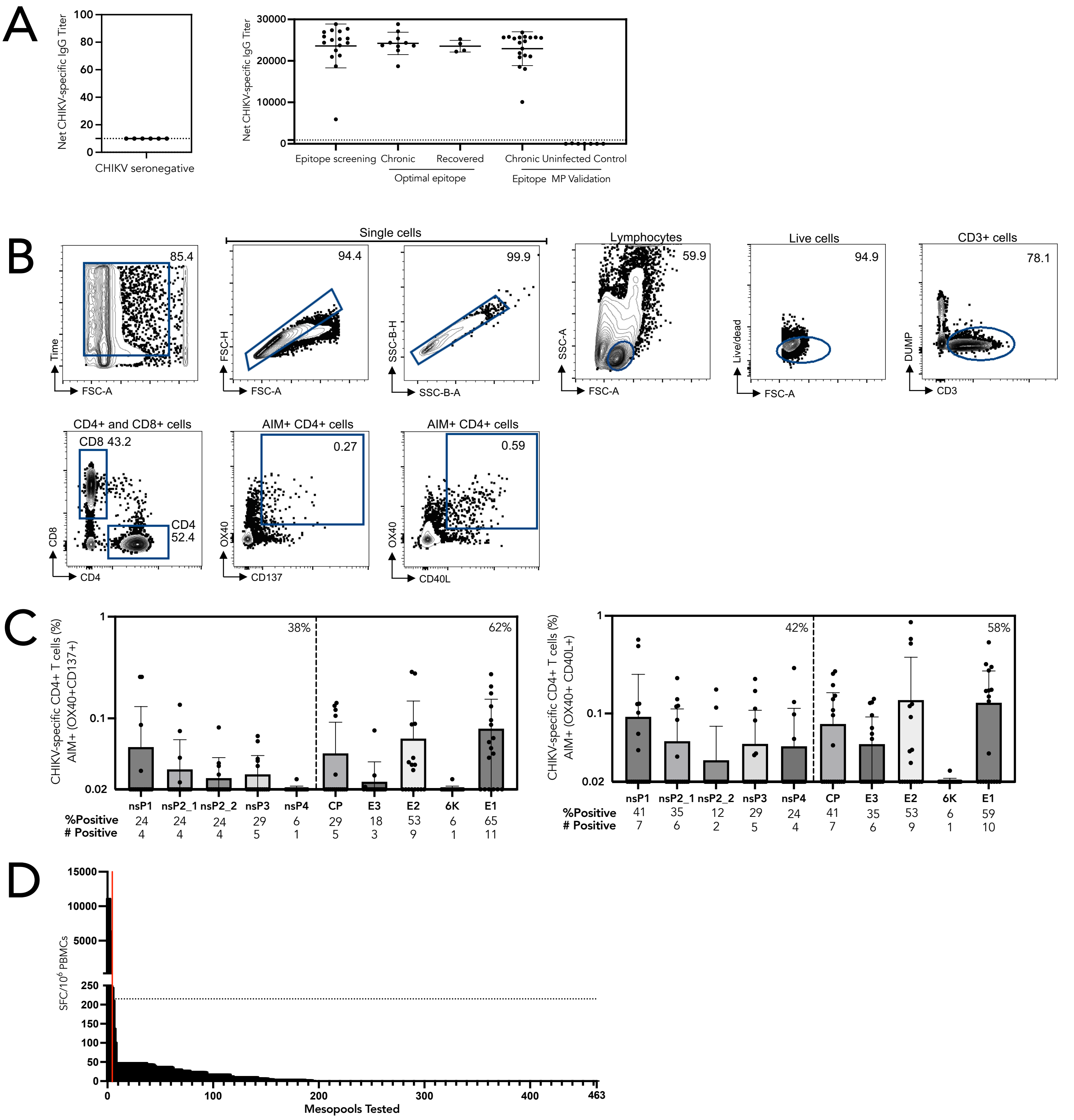


**Figure S1: Cohort serology, representative gating strategy, AIM+ CD4⁺ T cell responses and FluoroSpot Assay threshold of positivity**
**(A)** Net CHIKV-specific IgG titers measured in the study cohort. Left panel shows titers for samples collected from Nicaragua (threshold - 10). Right panel shows titers for samples collected from Colombia (threshold – 907). The threshold (dotted line) was used to determine seropositivity and to confirm CHIKV infection. **(B)** Representative gating strategy to define antigen-specific CD3+CD4⁺ cells by the AIM assay. The cells shown here were stimulated with nsP1 protein. **(C)** Antigen-specific CD4+ T cells quantified by AIM (left: OX40+CD137+ and right: OX40+CD40L+) after 24-hour stimulation with all CHIKV MPs in 17 CHIKV seropositive donors with chronic CHIKVD. The dotted line represents the limit of sensitivity (LOS; 0.02%). Data are represented as geomean ± geometric SD. **(D)** Graph shows the IFN$\gamma$ producing SFC/$10^6$ PBMCs for six CHIKV-seronegative donors post *in vitro* stimulation with all CHIKV mesopools. Responses were plotted in descending order of magnitude of response and the top 1% response (red line) was calculated. 99% of the response was lower than 215 SFC/$10^6$ PBMCs (dotted line), which was used as a threshold of positivity for following experiments

### FIGURE S2

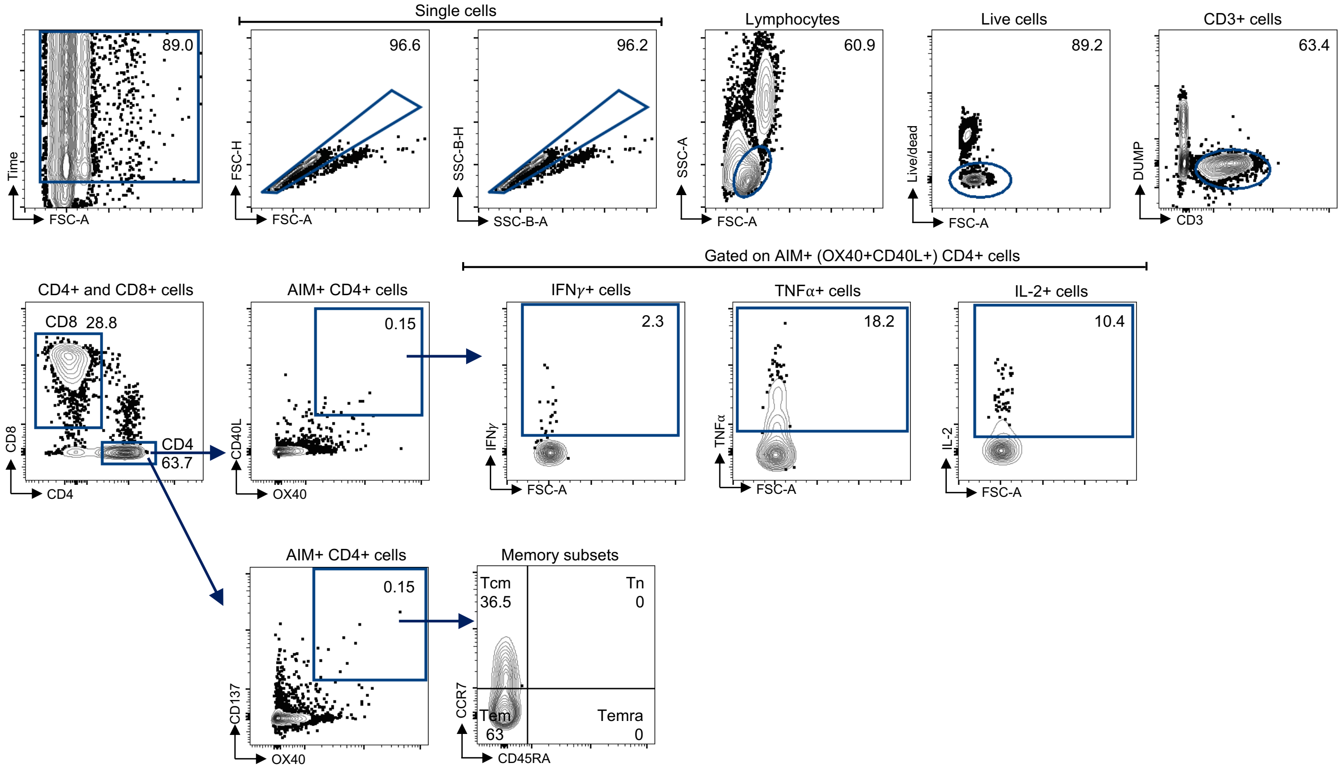


**Figure S2: Representative gating strategy for the AIM/ICS assay**
Representative gating strategy to define antigen-specific CD4+ T cells (OX40+CD137 and Ox40+CD40L+) and their functional profile via ICS staining of cytokines (IFNγ, TNFα and IL-2). Cells shown here were stimulated with the combined CHIKV S+NS epitope MP for initial gating and AIM gating, except with PHA for the cytokines.

### Table S1: Variant epitopes

| | Ref Sequence | Variant Epitope | Megapool | Screened in |
|---|---|---|---|---|
| 1 | PRGAIKVTAQPTDHV | PRGAIKVTAQLTDHV | nsP2_2 | 1 / 2 |
| 2 | KVTAQPTDHVVGEYL | KVTAQLTDHVVGEYL | nsP2_2 | 1 / 2 |
| 3 | PTDHVVGEYLVLSPQ | LTDHVVGEYLVLSPQ | nsP2_2 | 1 / 2 |
| 4 | FSNYSESEGDRELAA | FSNYTESEGDRELAA | nsP3 | 1 / 1 |
| 5 | HPPISFGASSETFPI | HPPISFGAPSETFPI | nsP3 | 1 / 1 |
| 6 | NEGEIESLSSELLTF | NDGEIESLSSELLTF | nsP3 | 1 / 1 |
| 7 | NRRYQPRPWTPRPTI | NRRYQPRPWTPRSTI | CP | 1 / 4 |
| 8 | RPRPQRQAGQLAQLI | RPRPQRKAGQLAQLI | CP | 1 / 4 |
| 9 | LAQLISAVNKLTMRA | LAQLISAVNKLTMRV | CP | 4 / 4 |
| 10 | SAVNKLTMRAVPQQK | SAVNKLTMRVVPQQK | CP | 2 / 4 |
| 11 | KPAQKKKKPGRRERM | KPVQKKKKPGRRERM | CP | 1 / 1 |
| 12 | DNVMRPGYYQLLQAS | DNVMSPGYYQLLQAS | E3 | 1 / 1 |
| 13 | MSQQSGNVKITVNGQ | MSQQSGNVKITVNSQ | E2 | 1 / 3 |
| 14 | GNVKITVNGQTVRYK | GNVKITVNSQTVRYK | E2 | 3 / 3 |
| 15 | TTTDKVINNCKVDQC | ITTDKVINNCKIDQC | E2 | 1 / 1 |
| 16 | KVDQCHAAVTNHKKW | KIDQCHAAVTNHKKW | E2 | 1 / 1 |
| 17 | VMHKKEVVLTVPTEG | VTHKKEIRLTVPTEG | E2 | 1 / 1 |
| 18 | EVVLTVPTEGLEVTW | EIRLTVPTEGLEVTW | E2 | 2 / 2 |
| 19 | LLSLICCIRTAKAAT | LLSLLCCIRTAKAAT | E2 | 1 / 2 |
| 20 | EQQPLFWLQALIPLA | EQQPLFWMQALIPLA | 6K | 2 / 3 |
| 21 | TQLVLQRPAVGTVHV | TQLVLQRPSAGTVHV | E1 | 1 / 1 |
| 22 | ATNPVRAVNCAVGNM | ATNPVRAMNCAVGNM | E1 | 3 / 5 |
| 23 | AVGNMPISIDIPEAA | AVGNMPISIDIPDAA | E1 | 2 / 4 |
| 24 | PISIDIPEAAFTRVV | PISIDIPDAAFTRVV | E1 | 2 / 4 |

### Table S2A: HLA binding predictions for N-term peptide in the immunodominant nsP1 region

| # | Allele | Core Sequence | Peptide Sequence | Percentile Rank |
|---|---|---|---|---|
| 1 | HLA-DPA1*02:01/DPB1*14:01 | GRRGKLSIM | DLTEGRRGKLSIMRG | 30 |
| 2 | HLA-DPA1*02:01/DPB1*05:01 | RRGKLSIMR | DLTEGRRGKLSIMRG | 53 |
| 3 | HLA-DRB1*11:01 | RRGKLSIMR | DLTEGRRGKLSIMRG | 64 |
| 4 | HLA-DRB1*08:02 | RRGKLSIMR | DLTEGRRGKLSIMRG | 70 |
| 5 | HLA-DRB5*01:01 | RRGKLSIMR | DLTEGRRGKLSIMRG | 71 |
| 6 | HLA-DPA1*01:03/DPB1*02:01 | RRGKLSIMR | DLTEGRRGKLSIMRG | 73 |
| 7 | HLA-DRB1*13:02 | RRGKLSIMR | DLTEGRRGKLSIMRG | 74 |
| 8 | HLA-DQA1*05:01/DQB1*03:01 | GRRGKLSIM | DLTEGRRGKLSIMRG | 77 |
| 9 | HLA-DQA1*04:01/DQB1*04:02 | TEGRRGKLS | DLTEGRRGKLSIMRG | 81 |
| 10 | HLA-DRB4*01:01 | RRGKLSIMR | DLTEGRRGKLSIMRG | 83 |
| 11 | HLA-DRB3*02:02 | RRGKLSIMR | DLTEGRRGKLSIMRG | 86 |
| 12 | HLA-DPA1*02:01/DPB1*01:01 | RRGKLSIMR | DLTEGRRGKLSIMRG | 87 |
| 13 | HLA-DPA1*03:01/DPB1*04:02 | RRGKLSIMR | DLTEGRRGKLSIMRG | 87 |
| 14 | HLA-DQA1*01:02/DQB1*06:02 | GRRGKLSIM | DLTEGRRGKLSIMRG | 87 |
| 15 | HLA-DRB1*03:01 | RRGKLSIMR | DLTEGRRGKLSIMRG | 89 |
| 16 | HLA-DRB1*01:01 | RGKLSIMRG | DLTEGRRGKLSIMRG | 91 |
| 17 | HLA-DPA1*01:03/DPB1*04:01 | RRGKLSIMR | DLTEGRRGKLSIMRG | 92 |
| 18 | HLA-DRB1*12:01 | TEGRRGKLS | DLTEGRRGKLSIMRG | 92 |
| 19 | HLA-DRB1*15:01 | RGKLSIMRG | DLTEGRRGKLSIMRG | 94 |
| 20 | HLA-DRB1*09:01 | GRRGKLSIM | DLTEGRRGKLSIMRG | 94 |
| 21 | HLA-DQA1*03:01/DQB1*03:02 | GRRGKLSIM | DLTEGRRGKLSIMRG | 95 |
| 22 | HLA-DRB1*04:01 | RRGKLSIMR | DLTEGRRGKLSIMRG | 97 |
| 23 | HLA-DQA1*05:01/DQB1*02:01 | GRRGKLSIM | DLTEGRRGKLSIMRG | 97 |
| 24 | HLA-DQA1*01:01/DQB1*05:01 | RGKLSIMRG | DLTEGRRGKLSIMRG | 98 |
| 25 | HLA-DRB1*04:05 | RRGKLSIMR | DLTEGRRGKLSIMRG | 98 |
| 26 | HLA-DRB1*07:01 | GRRGKLSIM | DLTEGRRGKLSIMRG | 98 |
| 27 | HLA-DRB3*01:01 | RRGKLSIMR | DLTEGRRGKLSIMRG | 99 |

### Table S2B: HLA binding predictions for C-term peptide in the immunodominant nsP1 region

| # | Allele | Core Sequence | Peptide Sequence | Percentile Rank |
|---|---|---|---|---|
| 1 | HLA-DRB1*03:01 | LKPCDRVLF | KKLKPCDRVLFSVGS | 11 |
| 2 | HLA-DRB1*12:01 | LKPCDRVLF | KKLKPCDRVLFSVGS | 14 |
| 3 | HLA-DRB1*11:01 | LKPCDRVLF | KKLKPCDRVLFSVGS | 26 |
| 4 | HLA-DRB1*13:02 | LKPCDRVLF | KKLKPCDRVLFSVGS | 28 |
| 5 | HLA-DRB3*01:01 | LKPCDRVLF | KKLKPCDRVLFSVGS | 47 |
| 6 | HLA-DRB1*08:02 | LKPCDRVLF | KKLKPCDRVLFSVGS | 59 |
| 7 | HLA-DRB1*15:01 | LKPCDRVLF | KKLKPCDRVLFSVGS | 68 |
| 8 | HLA-DPA1*02:01/DPB1*05:01 | LKPCDRVLF | KKLKPCDRVLFSVGS | 69 |
| 9 | HLA-DRB4*01:01 | LKPCDRVLF | KKLKPCDRVLFSVGS | 70 |
| 10 | HLA-DRB3*02:02 | LKPCDRVLF | KKLKPCDRVLFSVGS | 77 |
| 11 | HLA-DRB5*01:01 | LKPCDRVLF | KKLKPCDRVLFSVGS | 77 |
| 12 | HLA-DPA1*01:03/DPB1*02:01 | LKPCDRVLF | KKLKPCDRVLFSVGS | 81 |
| 13 | HLA-DPA1*02:01/DPB1*01:01 | LKPCDRVLF | KKLKPCDRVLFSVGS | 83 |
| 14 | HLA-DPA1*01:03/DPB1*04:01 | LKPCDRVLF | KKLKPCDRVLFSVGS | 84 |
| 15 | HLA-DPA1*03:01/DPB1*04:02 | LKPCDRVLF | KKLKPCDRVLFSVGS | 85 |
| 16 | HLA-DPA1*02:01/DPB1*14:01 | LKPCDRVLF | KKLKPCDRVLFSVGS | 87 |
| 17 | HLA-DRB1*09:01 | LKPCDRVLF | KKLKPCDRVLFSVGS | 87 |
| 18 | HLA-DRB1*07:01 | LKPCDRVLF | KKLKPCDRVLFSVGS | 88 |
| 19 | HLA-DQA1*01:01/DQB1*05:01 | LKPCDRVLF | KKLKPCDRVLFSVGS | 89 |
| 20 | HLA-DRB1*01:01 | LKPCDRVLF | KKLKPCDRVLFSVGS | 92 |
| 21 | HLA-DRB1*04:01 | LKPCDRVLF | KKLKPCDRVLFSVGS | 94 |
| 22 | HLA-DQA1*03:01/DQB1*03:02 | PCDRVLFSV | KKLKPCDRVLFSVGS | 95 |
| 23 | HLA-DRB1*04:05 | LKPCDRVLF | KKLKPCDRVLFSVGS | 95 |
| 24 | HLA-DQA1*04:01/DQB1*04:02 | LKPCDRVLF | KKLKPCDRVLFSVGS | 95 |
| 25 | HLA-DQA1*05:01/DQB1*02:01 | KPCDRVLFS | KKLKPCDRVLFSVGS | 97 |
| 26 | HLA-DQA1*01:02/DQB1*06:02 | LKPCDRVLF | KKLKPCDRVLFSVGS | 98 |
| 27 | HLA-DQA1*05:01/DQB1*03:01 | KPCDRVLFS | KKLKPCDRVLFSVGS | 99 |

### Table S3A: HLA binding predictions for N-term peptide in the immunodominant E1 region

| # | Allele | Core Sequence | Peptide Sequence | Percentile Rank |
|---|---|---|---|---|
| 1 | HLA-DQA1*05:01/DQB1*03:01 | SDFGGVAII | THSSDFGGVAIIKYA | 14 |
| 2 | HLA-DRB1*09:01 | FGGVAIIKY | THSSDFGGVAIIKYA | 19 |
| 3 | HLA-DPA1*03:01/DPB1*04:02 | FGGVAIIKY | THSSDFGGVAIIKYA | 20 |
| 4 | HLA-DQA1*03:01/DQB1*03:02 | FGGVAIIKY | THSSDFGGVAIIKYA | 21 |
| 5 | HLA-DPA1*02:01/DPB1*01:01 | FGGVAIIKY | THSSDFGGVAIIKYA | 22 |
| 6 | HLA-DRB1*07:01 | FGGVAIIKY | THSSDFGGVAIIKYA | 23 |
| 7 | HLA-DPA1*01:03/DPB1*04:01 | FGGVAIIKY | THSSDFGGVAIIKYA | 26 |
| 8 | HLA-DQA1*01:02/DQB1*06:02 | FGGVAIIKY | THSSDFGGVAIIKYA | 27 |
| 9 | HLA-DQA1*05:01/DQB1*02:01 | FGGVAIIKY | THSSDFGGVAIIKYA | 28 |
| 10 | HLA-DRB1*12:01 | FGGVAIIKY | THSSDFGGVAIIKYA | 28 |
| 11 | HLA-DRB3*01:01 | FGGVAIIKY | THSSDFGGVAIIKYA | 40 |
| 12 | HLA-DPA1*02:01/DPB1*05:01 | FGGVAIIKY | THSSDFGGVAIIKYA | 44 |
| 13 | HLA-DRB5*01:01 | FGGVAIIKY | THSSDFGGVAIIKYA | 47 |
| 14 | HLA-DRB1*04:05 | FGGVAIIKY | THSSDFGGVAIIKYA | 47 |
| 15 | HLA-DPA1*01:03/DPB1*02:01 | FGGVAIIKY | THSSDFGGVAIIKYA | 48 |
| 16 | HLA-DPA1*02:01/DPB1*14:01 | FGGVAIIKY | THSSDFGGVAIIKYA | 54 |
| 17 | HLA-DQA1*01:01/DQB1*05:01 | FGGVAIIKY | THSSDFGGVAIIKYA | 56 |
| 18 | HLA-DRB1*04:01 | FGGVAIIKY | THSSDFGGVAIIKYA | 56 |
| 19 | HLA-DRB1*08:02 | FGGVAIIKY | THSSDFGGVAIIKYA | 57 |
| 20 | HLA-DQA1*04:01/DQB1*04:02 | FGGVAIIKY | THSSDFGGVAIIKYA | 59 |
| 21 | HLA-DRB3*02:02 | FGGVAIIKY | THSSDFGGVAIIKYA | 63 |
| 22 | HLA-DRB1*13:02 | FGGVAIIKY | THSSDFGGVAIIKYA | 64 |
| 23 | HLA-DRB1*11:01 | FGGVAIIKY | THSSDFGGVAIIKYA | 66 |
| 24 | HLA-DRB1*01:01 | FGGVAIIKY | THSSDFGGVAIIKYA | 68 |
| 25 | HLA-DRB1*03:01 | FGGVAIIKY | THSSDFGGVAIIKYA | 78 |
| 26 | HLA-DRB1*15:01 | FGGVAIIKY | THSSDFGGVAIIKYA | 78 |
| 27 | HLA-DRB4*01:01 | FGGVAIIKY | THSSDFGGVAIIKYA | 79 |

### Table S3B: HLA binding predictions for C-term peptide in the immunodominant E1 region

| # | Allele | Core Sequence | Peptide Sequence | Percentile Rank |
|---|---|---|---|---|
| 1 | HLA-DPA1*02:01/DPB1*14:01 | KCAVHSMTN | ASKKGKCAVHSMTNA | 36 |
| 2 | HLA-DQA1*04:01/DQB1*04:02 | KGKCAVHSM | ASKKGKCAVHSMTNA | 68 |
| 3 | HLA-DQA1*01:02/DQB1*06:02 | KGKCAVHSM | ASKKGKCAVHSMTNA | 78 |
| 4 | HLA-DPA1*02:01/DPB1*05:01 | KKGKCAVHS | ASKKGKCAVHSMTNA | 79 |
| 5 | HLA-DRB1*08:02 | KKGKCAVHS | ASKKGKCAVHSMTNA | 82 |
| 6 | HLA-DRB1*11:01 | KKGKCAVHS | ASKKGKCAVHSMTNA | 83 |
| 7 | HLA-DPA1*01:03/DPB1*02:01 | KKGKCAVHS | ASKKGKCAVHSMTNA | 83 |
| 8 | HLA-DQA1*05:01/DQB1*03:01 | SKKGKCAVH | ASKKGKCAVHSMTNA | 83 |
| 9 | HLA-DPA1*01:03/DPB1*04:01 | KKGKCAVHS | ASKKGKCAVHSMTNA | 84 |
| 10 | HLA-DRB1*04:01 | KCAVHSMTN | ASKKGKCAVHSMTNA | 86 |
| 11 | HLA-DRB1*07:01 | KGKCAVHSM | ASKKGKCAVHSMTNA | 86 |
| 12 | HLA-DRB5*01:01 | KGKCAVHSM | ASKKGKCAVHSMTNA | 88 |
| 13 | HLA-DPA1*02:01/DPB1*01:01 | KGKCAVHSM | ASKKGKCAVHSMTNA | 89 |
| 14 | HLA-DRB3*02:02 | GKCAVHSMT | ASKKGKCAVHSMTNA | 91 |
| 15 | HLA-DRB4*01:01 | KCAVHSMTN | ASKKGKCAVHSMTNA | 91 |
| 16 | HLA-DPA1*03:01/DPB1*04:02 | KKGKCAVHS | ASKKGKCAVHSMTNA | 91 |
| 17 | HLA-DRB1*01:01 | KCAVHSMTN | ASKKGKCAVHSMTNA | 92 |
| 18 | HLA-DRB1*04:05 | KCAVHSMTN | ASKKGKCAVHSMTNA | 93 |
| 19 | HLA-DRB1*13:02 | KKGKCAVHS | ASKKGKCAVHSMTNA | 94 |
| 20 | HLA-DRB1*09:01 | GKCAVHSMT | ASKKGKCAVHSMTNA | 94 |
| 21 | HLA-DRB1*03:01 | GKCAVHSMT | ASKKGKCAVHSMTNA | 95 |
| 22 | HLA-DQA1*03:01/DQB1*03:02 | KGKCAVHSM | ASKKGKCAVHSMTNA | 96 |
| 23 | HLA-DQA1*01:01/DQB1*05:01 | KGKCAVHSM | ASKKGKCAVHSMTNA | 96 |
| 24 | HLA-DRB1*15:01 | KCAVHSMTN | ASKKGKCAVHSMTNA | 97 |
| 25 | HLA-DRB1*12:01 | KGKCAVHSM | ASKKGKCAVHSMTNA | 97 |
| 26 | HLA-DQA1*05:01/DQB1*02:01 | KGKCAVHSM | ASKKGKCAVHSMTNA | 98 |
| 27 | HLA-DRB3*01:01 | KCAVHSMTN | ASKKGKCAVHSMTNA | 99 |

#### Table S4A: HLA binding predictions for N-term peptide in the immunodominant CP region

| # | Allele | Core Sequence | Peptide Sequence | Percentile Rank |
|---|---|---|---|---|
| 1 | HLA-DQA1*01:02/DQB1*06:02 | GQLAQLISA | RQAGQLAQLISAVNK | 11 |
| 2 | HLA-DRB1*01:01 | LAQLISAVN | RQAGQLAQLISAVNK | 16 |
| 3 | HLA-DRB1*04:05 | LAQLISAVN | RQAGQLAQLISAVNK | 17 |
| 4 | HLA-DRB1*04:01 | LAQLISAVN | RQAGQLAQLISAVNK | 23 |
| 5 | HLA-DPA1*02:01/DPB1*14:01 | LAQLISAVN | RQAGQLAQLISAVNK | 25 |
| 6 | HLA-DRB4*01:01 | LAQLISAVN | RQAGQLAQLISAVNK | 27 |
| 7 | HLA-DRB1*08:02 | LAQLISAVN | RQAGQLAQLISAVNK | 31 |
| 8 | HLA-DRB1*12:01 | LAQLISAVN | RQAGQLAQLISAVNK | 33 |
| 9 | HLA-DRB3*02:02 | LAQLISAVN | RQAGQLAQLISAVNK | 37 |
| 10 | HLA-DRB5*01:01 | LAQLISAVN | RQAGQLAQLISAVNK | 37 |
| 11 | HLA-DRB1*11:01 | LAQLISAVN | RQAGQLAQLISAVNK | 39 |
| 12 | HLA-DRB1*15:01 | LAQLISAVN | RQAGQLAQLISAVNK | 41 |
| 13 | HLA-DPA1*03:01/DPB1*04:02 | LAQLISAVN | RQAGQLAQLISAVNK | 42 |
| 14 | HLA-DQA1*05:01/DQB1*02:01 | QLAQLISAV | RQAGQLAQLISAVNK | 47 |
| 15 | HLA-DQA1*04:01/DQB1*04:02 | QLAQLISAV | RQAGQLAQLISAVNK | 47 |
| 16 | HLA-DQA1*03:01/DQB1*03:02 | LAQLISAVN | RQAGQLAQLISAVNK | 48 |
| 17 | HLA-DQA1*05:01/DQB1*03:01 | GQLAQLISA | RQAGQLAQLISAVNK | 51 |
| 18 | HLA-DRB1*09:01 | LAQLISAVN | RQAGQLAQLISAVNK | 51 |
| 19 | HLA-DPA1*02:01/DPB1*01:01 | LAQLISAVN | RQAGQLAQLISAVNK | 52 |
| 20 | HLA-DPA1*02:01/DPB1*05:01 | LAQLISAVN | RQAGQLAQLISAVNK | 53 |
| 21 | HLA-DQA1*01:01/DQB1*05:01 | LAQLISAVN | RQAGQLAQLISAVNK | 55 |
| 22 | HLA-DPA1*01:03/DPB1*02:01 | LAQLISAVN | RQAGQLAQLISAVNK | 59 |
| 23 | HLA-DRB1*07:01 | LAQLISAVN | RQAGQLAQLISAVNK | 59 |
| 24 | HLA-DPA1*01:03/DPB1*04:01 | LAQLISAVN | RQAGQLAQLISAVNK | 63 |
| 25 | HLA-DRB1*13:02 | LAQLISAVN | RQAGQLAQLISAVNK | 76 |
| 26 | HLA-DRB1*03:01 | LAQLISAVN | RQAGQLAQLISAVNK | 80 |
| 27 | HLA-DRB3*01:01 | LAQLISAVN | RQAGQLAQLISAVNK | 84 |

#### Table S4B: HLA binding predictions for C-term peptide in the immunodominant CP region

| # | Allele | Core Sequence | Peptide Sequence | Percentile Rank |
|---|---|---|---|---|
| 1 | HLA-DRB1*03:01 | AVPQQKPRR | LTMRAVPQQKPRRNR | 12 |
| 2 | HLA-DRB1*13:02 | AVPQQKPRR | LTMRAVPQQKPRRNR | 21 |
| 3 | HLA-DRB4*01:01 | AVPQQKPRR | LTMRAVPQQKPRRNR | 25 |
| 4 | HLA-DRB1*11:01 | AVPQQKPRR | LTMRAVPQQKPRRNR | 35 |
| 5 | HLA-DRB1*12:01 | MRAVPQQKP | LTMRAVPQQKPRRNR | 36 |
| 6 | HLA-DRB1*08:02 | MRAVPQQKP | LTMRAVPQQKPRRNR | 39 |
| 7 | HLA-DRB5*01:01 | MRAVPQQKP | LTMRAVPQQKPRRNR | 41 |
| 8 | HLA-DPA1*02:01/DPB1*05:01 | AVPQQKPRR | LTMRAVPQQKPRRNR | 41 |
| 9 | HLA-DPA1*02:01/DPB1*14:01 | RAVPQQKPR | LTMRAVPQQKPRRNR | 48 |
| 10 | HLA-DRB1*15:01 | MRAVPQQKP | LTMRAVPQQKPRRNR | 49 |
| 11 | HLA-DRB3*02:02 | AVPQQKPRR | LTMRAVPQQKPRRNR | 50 |
| 12 | HLA-DRB1*09:01 | MRAVPQQKP | LTMRAVPQQKPRRNR | 50 |
| 13 | HLA-DRB1*04:01 | MRAVPQQKP | LTMRAVPQQKPRRNR | 53 |
| 14 | HLA-DPA1*02:01/DPB1*01:01 | MRAVPQQKP | LTMRAVPQQKPRRNR | 54 |
| 15 | HLA-DQA1*01:02/DQB1*06:02 | MRAVPQQKP | LTMRAVPQQKPRRNR | 62 |
| 16 | HLA-DRB3*01:01 | AVPQQKPRR | LTMRAVPQQKPRRNR | 63 |
| 17 | HLA-DQA1*04:01/DQB1*04:02 | MRAVPQQKP | LTMRAVPQQKPRRNR | 64 |
| 18 | HLA-DPA1*03:01/DPB1*04:02 | MRAVPQQKP | LTMRAVPQQKPRRNR | 66 |
| 19 | HLA-DPA1*01:03/DPB1*04:01 | MRAVPQQKP | LTMRAVPQQKPRRNR | 67 |
| 20 | HLA-DPA1*01:03/DPB1*02:01 | MRAVPQQKP | LTMRAVPQQKPRRNR | 68 |
| 21 | HLA-DRB1*04:05 | MRAVPQQKP | LTMRAVPQQKPRRNR | 71 |
| 22 | HLA-DRB1*07:01 | MRAVPQQKP | LTMRAVPQQKPRRNR | 72 |
| 23 | HLA-DRB1*01:01 | MRAVPQQKP | LTMRAVPQQKPRRNR | 73 |
| 24 | HLA-DQA1*05:01/DQB1*03:01 | TMRAVPQQK | LTMRAVPQQKPRRNR | 78 |
| 25 | HLA-DQA1*03:01/DQB1*03:02 | MRAVPQQKP | LTMRAVPQQKPRRNR | 88 |
| 26 | HLA-DQA1*05:01/DQB1*02:01 | MRAVPQQKP | LTMRAVPQQKPRRNR | 90 |
| 27 | HLA-DQA1*01:01/DQB1*05:01 | MRAVPQQKP | LTMRAVPQQKPRRNR | 91 |

### Table S5: HLA alleles of fine epitope mapping cohort

| Donor ID | Disease status | Tested against | HLA-DPB1 Allele 1 | HLA-DPB1 Allele 2 | HLA-DQA1 Allele 1 | HLA-DQA1 Allele 2 | HLA-DQB1 Allele 1 | HLA-DQB1 Allele 2 | HLA-DRB1 Allele 1 | HLA-DRB1 Allele 2 | HLA-DRB3 Allele 1 | HLA-DRB3 Allele 2 | HLA-DRB4 Allele 1 | HLA-DRB4 Allele 2 | HLA-DRB5 Allele 1 | HLA-DRB5 Allele 2 |
|---|---|---|---|---|---|---|---|---|---|---|---|---|---|---|---|---|
| 1 | Chronic | nsP1 | DPB1*04:01 | DPB1*18:01 | DQA1*01:02 | DQA1*05:01 | DQB1*03:01 | DQB1*06:02 | DRB1*14:06 | DRB1*15:03 | DRB3*01:01 | 0 | 0 | 0 | DRB5*01:01 | 0 |
| 2 | Chronic | nsP1 | DPB1*04:01 | DPB1*04:02 | DQA1*01:01 | DQA1*05:01 | DQB1*03:01 | DQB1*05:01 | DRB1*01:01 | DRB1*11:01 | DRB3*02:02 | 0 | 0 | 0 | 0 | 0 |
| 3 | Chronic | nsP1, E1 | DPB1*04:02 | DPB1*18:01 | DQA1*01:02 | DQA1*03:01 | DQB1*03:02 | DQB1*06:02 | DRB1*04:11 | DRB1*15:03 | 0 | 0 | DRB4*01:01 | 0 | DRB5*01:01 | 0 |
| 4 | Chronic | E1 | DPB1*04:02 | DPB1*05:01 | DQA1*01:02 | DQA1*03:01 | DQB1*03:02 | DQB1*06:09 | DRB1*04:07 | DRB1*13:02 | DRB3*03:01 | 0 | DRB4*01:01 | 0 | 0 | 0 |
| 5 | Chronic | E1, CP | DPB1*06:01 | DPB1*10:01 | DQA1*01:03 | DQA1*04:01 | DQB1*04:02 | DQB1*06:03 | DRB1*08:02 | DRB1*13:01 | DRB3*01:01 | 0 | 0 | 0 | 0 | 0 |
| 6 | Chronic | E1 | DPB1*04:01 | DPB1*04:01 | DQA1*04:01 | DQA1*05:01 | DQB1*03:01 | DQB1*04:02 | DRB1*08:02 | DRB1*13:03 | DRB3*01:01 | 0 | 0 | 0 | 0 | 0 |
| 7 | Chronic | E1 | DPB1*04:02 | DPB1*14:01 | DQA1*03:01 | DQA1*03:01 | DQB1*03:02 | DQB1*04:02 | DRB1*04:11 | DRB1*04:11 | 0 | 0 | DRB4*01:01 | DRB4*01:01 | 0 | 0 |
| 8 | Chronic | CP | DPB1*04:01 | DPB1*04:01 | DQA1*04:01 | DQA1*05:01 | DQB1*02:01 | DQB1*04:02 | DRB1*03:01 | DRB1*08:02 | DRB3*01:01 | 0 | 0 | 0 | 0 | 0 |
| 9 | Chronic | CP | DPB1*04:01 | DPB1*04:02 | DQA1*03:01 | DQA1*05:01 | DQB1*02:01 | DQB1*03:02 | DRB1*03:01 | DRB1*04:11 | DRB3*02:02 | 0 | DRB4*01:01 | 0 | 0 | 0 |
| 10 | Chronic | CP | DPB1*03:01 | DPB1*03:01 | DQA1*01:02 | DQA1*05:01 | DQB1*03:01 | DQB1*06:04 | DRB1*03:02 | DRB1*13:02 | DRB3*01:01 | DRB3*03:01 | 0 | 0 | 0 | 0 |
| 11 | Recovered | nsP1 | DPB1*03:01 | DPB1*04:02 | DQA1*01:02 | DQA1*05:01 | DQB1*03:01 | DQB1*06:04 | DRB1*08:04 | DRB1*13:02 | DRB3*03:01 | 0 | 0 | 0 | 0 | 0 |
| 12 | Recovered | nsP1, E1 | DPB1*04:01 | DPB1*04:02 | DQA1*01:02 | DQA1*05:01 | DQB1*03:01 | DQB1*05:02 | DRB1*11:03 | DRB1*15:01 | DRB3*02:02 | 0 | 0 | 0 | DRB5*01:01 | 0 |
| 13 | Recovered | E1 | DPB1*02:01 | DPB1*04:01 | DQA1*01:02 | DQA1*01:03 | DQB1*06:02 | DQB1*06:03 | DRB1*13:01 | DRB1*15:01 | DRB3*01:01 | 0 | 0 | 0 | DRB5*01:01 | 0 |
| 14 | Recovered | E1 | DPB1*04:01 | DPB1*17:01 | DQA1*02:01 | DQA1*03:01 | DQB1*02:02 | DQB1*03:02 | DRB1*04:02 | DRB1*07:01 | 0 | 0 | DRB4*01:01 | DRB4*01:01 | 0 | 0 |

### Table S6: Antibodies used in AIM assay

| Antibody | Clone (Vendor) | Catalog no. |
|---|---|---|
| CD40 | HB14 (Miltenyi Biotec) | 130-108-041 |
| CCR6-BUV496 | 11A9 (BD Biosciences) | 612948 |
| CXCR5-BV421 | J252D4 (BioLegend) | 356920 |
| CXCR3-BV605 | G025H7 (BioLegend) | 353728 |
| CCR7-BV711 | G043H7 (BioLegend) | 353228 |
| LIVE/DEAD-Fixable Blue | (ThermoFisher) | L23105 |
| CD3-BUV395 | UCHT1 (BD Biosciences) | 563546 |
| CD8-BUV805 | SK1 (BD Biosciences) | 612889 |
| CD16-BV510 | 3G8 (BioLegend) | 302048 |
| CD14-BV510 | 63D3 (BioLegend) | 367124 |
| CD20-BV510 | 2H7 (BioLegend) | 302340 |
| CD45RA-BV570 | Hl100 (BioLegend) | 304132 |
| CD4-cFluor b548 | SK3 (Cytek) | R7-20043 |
| CD95-BB700 | DX2 (BD Biosciences) | 566542 |
| HLA-DR-APC-R700 | G46-6 (BD Biosciences) | 565127 |
| CD38-BV650 | HB-7 (BioLegend) | 356620 |
| PD-1-BV785 | EH12.2H7 (BioLegend) | 329930 |
| CD40L-PE-Dazzle594 | 24-31 (BioLegend) | 310840 |
| OX40-APC | Ber-Act35 (BioLegend) | 350008 |
| CD69-FITC | FN50 (BioLegend) | 310904 |
| CD137-BUV737 | 4b4-1 (BD Biosciences) | 741861 |

### Table S7: Antibodies used in AIM/ICS assay

| Antibody | Clone (Source) | Catalog no. |
|---|---|---|
| CD40 | HB14 (Miltenyi Biotec) | 130-094-133 |
| CCR6-BUV496 | 11A9 (Biolegend) | 612948 |
| CXCR5-BV421 | J252D4 (Biolegend) | 356920 |
| CXCR3-BV605 | G025H7 (Biolegend) | 353728 |
| CCR7-BV711 | G043H7 (Biolegend) | 353228 |
| CCR4-APC | L291H4 (Biolegend) | 359408 |
| CD69 - PE | FN50 (BD Biosciences) | 555531 |
| CD137 - PE-Cy5 | 4B4-1 (BD Biosciences) | 551137 |
| LIVE/DEAD-Fixable Blue | (ThermoFisher) | L23105 |
| CD3-BUV395 | UCHT1 (BD Biosciences) | 563546 |
| CD8-BUV805 | SK1 (BD Biosciences) | 612889 |
| CD16-BV510 | 3G8 (Biolegend) | 302048 |
| CD14-BV510 | 63D3 (Biolegend) | 367124 |
| CD20-BV510 | 2H7 (Biolegend) | 302340 |
| CD45RA-BV570 | HI100 (Biolegend) | 304132 |
| CD4-cFluor b548 | SK3 (Cytek) | R7-20043 |
| HLA-DR-APC R700 | G46-6 (BD Biosciences) | 565127 |
| CD38-BV650 | HB-7 (Biolegend) | 353228 |
| PD-1-BV480 | EH12.1 (BD Biosciences) | 566112 |
| OX40-PE-Cy7 | Ber-ACT35 (Biolegend) | 350012 |
| IFNg-FITC | 4S.B3 (eBioscience) | 11-7319-82 |
| IL-17-BV785 | BL168 (Biolegend) | 512338 |
| IL-10-PE-Dazzle594 | JES3-19F1 (Biolegend) | 506812 |
| IL-2-BUV737 | (BD Biosciences) | 612836 |
| TNFa-eFluor450 | Mab11 (eBiosciences) | 48-7349-42 |
| Granzyme B-AF647 | GR11 (BD Biosciences) | 560212 |
| CD40L-APC-efluor 780 | 24-31 (ThermoFisher) | 46-1548-42 |
